# Molecular determinants of Neu5Ac binding to a tripartite ATP independent periplasmic (TRAP) transporter

**DOI:** 10.1101/2024.03.29.587382

**Authors:** Parveen Goyal, KanagaVijayan Dhanabalan, Mariafrancesca Scalise, Rosmarie Friemann, Cesare Indiveri, Renwick C.J. Dobson, Kutti R. Vinothkumar, Subramanian Ramaswamy

## Abstract

*N*-Acetylneuraminic acid (Neu5Ac) is a negatively charged nine-carbon amino-sugar that is often the peripheral sugar in human cell-surface glycoconjugates. Some bacteria scavenge, import, and metabolize Neu5Ac or redeploy it on their cell surfaces for immune evasion. The import of Neu5Ac by many bacteria is mediated by tripartite ATP-independent periplasmic (TRAP) transporters. We have previously reported the structures of SiaQM, a membrane-embedded component of the *Haemophilus influenzae* TRAP transport system, (Currie *et al.,* 2024). However, none of the published structures contain Neu5Ac bound to SiaQM. This information is critical for defining the transport mechanism and for further structure-activity relationship studies. Here, we report the structures of *Fusobacterium nucleatum* SiaQM with and without Neu5Ac. Both structures are in an inward (cytoplasmic side) facing conformation. The Neu5Ac-bound structure reveals the interactions of Neu5Ac with the transporter and its relationship with the Na^+^ binding sites. Two of the Na^+^-binding sites are similar to those described previously. We identify a third metal-binding site that is further away and buried in the elevator domain. Ser300 and Ser345 interact with the C1-carboxylate group of Neu5Ac. Proteoliposome-based transport assays showed that Ser300-Neu5Ac interaction is critical for transport, whereas Ser345 is dispensable. Neu5Ac primarily interacts with residues in the elevator domain of the protein, thereby supporting the elevator with an operator mechanism. The residues interacting with Neu5Ac are conserved, providing fundamental information required to design inhibitors against this class of proteins.

## Introduction

Sialic acids are nine-carbon amino sugars common on the surface of mammalian cells and on secreted molecules. *N*-Acetylneuraminic acid (Neu5Ac) and *N*-glycolylneuraminic acid (Neu5Gc) are the most prevalent forms in mammals (Lewis and Lewis, 2012). A dynamic interplay between genetics, environmental cues, and cell signaling orchestrates the intricate regulation of sialic acid synthesis in mammals. These sugars frequently occupy terminal positions in glycolipids and glycoproteins and play pivotal roles in cell-cell interactions such as signaling, adhesion, and recognition (Chen and Varki, 2010). Notably, a frameshift mutation disrupts the CMP-Neu5Ac hydrolase (CMAH) gene and leads to the loss of enzymatic activity, resulting in Neu5Ac being the sole outermost sugar in humans. Conversely, primates possessing a functional CMP-Neu5Ac hydrolase, Neu5Gc is typically found as the outermost sugar (Chou *et al*., 1998).

This evolutionary divergence has profound consequences for the interactions between humans and pathogenic agents, including bacteria, parasites, and viruses (Varki, 2009). While commensal bacteria harness Neu5Ac as a carbon source, pathogenic bacteria, such as *Haemophilus influenzae* (*Hi*) and *Fusobacterium nucleatum* (*Fn*), evolved the ability to add Neu5Ac as the outermost sugar in their cell surface glycol-conjugates and use molecular mimicry to evade the immune system (Bell *et al*., 2023; Severi *et al*., 2007). Sialic acid is abundant in many niches [*e.g.*, [Neu5Ac] in human serum is 1.6–2.2 mM (Sillanaukee *et al*., 1999)]. However, the concentration of free sialic acid is often low (approximately 0.2% of the total), and much of the sialic acid is conjugated to other macromolecules on the cell surface and is, therefore, not immediately available. Secreted sialidases from bacteria cleave Neu5Ac from mucins and other biomolecules within their niche (Sillanaukee *et al*., 1999). Cleaved Neu5Ac is transported into the periplasm of Gram-negative bacteria via porin-like *β-*barrel proteins. NanC from *E. coli* is the best-studied porin specific for sialic acids (Wirth *et al*., 2009).

Our groups have made significant strides in elucidating the structural intricacies of the proteins involved in sequestration, uptake, catabolism, and incorporation of Neu5Ac by bacteria (Bose *et al*., 2019; Coombes *et al*., 2020; Currie *et al*., 2021; Davies *et al*., 2019; Horne *et al*., 2020; Kumar *et al*., 2018; Manjunath *et al*., 2018; North *et al*., 2016; Setty *et al*., 2018). Bacteria rely on specialized Neu5Ac transporters within their cytosolic membranes, and four distinct classes of Neu5Ac transporters have been identified: sodium solute symporters (SSS) (North *et al*., 2018), major facilitator superfamily (MFS), ATP-binding cassette (ABC), and tripartite ATP-independent periplasmic (TRAP) transporters (Davies *et al*., 2023; Peter *et al*., 2022). TRAP transporters that manifest as two- or three-component systems consisting of a substrate-binding protein (SiaP) and transmembrane protein (SiaQM) (Rosa *et al*., 2018) are of particular interest due to their absence in eukaryotes. Deleting the sialic acid transporter abolishes Neu5Ac uptake, rendering bacteria incapable of incorporating Neu5Ac into lipopolysaccharides. Without a SiaQM transporter, bacteria form defective biofilms, have a lower cell density, and experience higher cell death (Allen *et al*., 2005). Additionally, amino acids, C4-dicarboxylates, aromatic substrates and alpha-keto acids are also transported by TRAP transporters (Vetting *et al*., 2015).

The first structures of SiaP were obtained from *H. influenzae* in an unliganded form and bound to Neu5Ac (Johnston *et al*., 2008; Müller *et al*., 2006). The binding of Neu5Ac to SiaP results in the closure of the two domains of the protein and the mechanism has been described as a “Venus fly trap” mechanism, a common phenomenon observed in many ligand-binding proteins (Felder *et al*., 1999). Structural and thermodynamic analyses of SiaP from *F. nucleatum*, *V. cholerae*, and *P. multocida* have revealed a conserved binding site, dissociation constant of Neu5Ac in the nanomolar range, and enthalpically driven substrate binding (Setty *et al*., 2018). Using smFRET on *Vc*SiaP, Peter and co-workers showed that conformational switching is strictly substrate-induced and that binding of the substrate stabilizes the interactions between the two domains (Peter *et al*., 2021). All the SiaP structures show the presence of a conserved Arginine that binds to the C1-carboxylate of Neu5Ac, and this Arg residue is critical as the high electrostatic affinity may be important to have a strong binding affinity that sequesters the small amounts that reach the bacterial periplasmic space (Glaenzer *et al*., 2017).

The second component of the TRAP transporter is the transmembrane protein SiaQM, which is characterized by two distinct domains: the Q-domain and M-domain. Two SiaQM transporter structures have been reported by cryo-EM, one from *H. influenzae* (*Hi*SiaQM) ranging from 2.95-4.70 Å resolution and another from *Protobacterium profundum* (*Pp*SiaQM) at 2.97 Å resolution (Currie *et al*., 2024; Davies *et al*., 2023; Peter *et al*., 2022).

The first structure of *Hi*SiaQM (4.7 Å resolution) demonstrated that it is composed of 15 transmembrane helices (TM) and two helical hairpins. *Hi*SiaQM likely functions as a monomeric unit, although a dimeric form has recently been reported (Currie *et al*., 2024; Peter *et al*., 2022). Similarly, *Pp*SiaQM is composed of 16 TM helices (Davies *et al*., 2023). Based on these structures, an elevator-type mechanism has been proposed for TRAP transporters to move substrates from the periplasm to the cytoplasm (Peter *et al*., 2024). Apart from the dimeric HiSiaQM structure, the other two structures determined used a megabody bound to SiaQM for cryo-EM analysis. In both cases, the megabody was bound to the extracellular side of the QM complex, featuring a deep open cavity on the cytosolic side (**Supplementary Figure 1**). This observation suggested that the transporter was captured in an inward-facing conformation, although the higher resolution dimeric HiSiaQM structure, even without a fiducial megabody, is also in the inward-facing conformation. Notably, the transport of Neu5Ac by TRAP transporters requires at least two sodium ions (Currie *et al*., 2024; Davies *et al*., 2023; Mulligan *et al*., 2012, 2009).

In sialic acid-rich environments such as the gut and saliva, bacterial virulence often correlates with their capacity to utilize Neu5Ac as a carbon source (Almagro-Moreno and Boyd, 2010; Corfield, 2015; Haines-Menges *et al*., 2015). Although inhibitors targeting viral sialidases (neuraminidases) have been developed as antiviral agents (*e*.*g*., oseltamivir, zanamivir, and peramivir) (Glanz *et al*., 2018), no drug currently targets bacterial infections by inhibiting sialic acid sequestration, uptake, catabolism, or incorporation. Human analogs of TRAP transporters are notably absent, underscoring their potential as promising therapeutic targets for combatting bacterial infections.

Both *Hi*SiaQM and *Pp*SiaQM structures lack information on the Neu5Ac binding site, which was identified based on modeling studies that relied on the ligand-bound structure of *Vc*INDY (Kinz-Thompson *et al*., 2022). Moreover, only the structures of SiaQM in the elevator-down conformation (inward-facing) have been reported, and further conformations along the transport cycle remains to be elucidated. The conformation of Neu5Ac bound to the transport domain may provide clues as to how it is received from the substrate-binding protein. In this study, we present the cryo-EM structures of SiaQM from *F. nucleatum*, both in its unliganded form and Neu5Ac bound form. The unliganded structure has density for two Na^+^-binding sites, whereas the liganded form has density for two Na^+^-binding sites and an additional metal-binding site.

## Results and Discussion

### Construct design, isolation, and strategy for structural determination

The TRAP transporter from *F. nucleatum* (*Fn*SiaQM) was tagged with green fluorescent protein (GFP) at the C-terminus and expressed in BL21 (DE3) cells for protein expression and detergent stabilization trials. Detergent scouting was performed to identify the preferred detergent for *Fn*SiaQM purification using fluorescence detection size-exclusion chromatography (Kawate and Gouaux, 2006). n-Dodecyl-β-D-maltopyranoside (DDM) was chosen as it solubilized and stabilized the protein better than other detergents. *Fn*SiaQM was re-cloned into the pBAD vector with N-terminal 6X -histidine followed by Strep-tag II for large-scale purification and expressed in TOP10 (Invitrogen) cells. The purified *Fn*SiaQM protein in the detergent micelles eluted as a monodisperse peak in size exclusion chromatography (**Supplementary Figure 2**). Subsequently, *Fn*SiaQM was further reconstituted in MSP1D1 nanodiscs, along with *E. coli* polar lipids.

Initially, we attempted to determine the *Fn*SiaQM structure using X-ray crystallography. After failing to obtain well-diffracting crystals, we switched to single-particle electron cryo-microscopy (cryo-EM). To achieve a reasonable size and better particle alignment, we raised nanobodies against *Fn*SiaQM in Alpaca (Center for Molecular Medicine, University of Kentucky College of Medicine). Two high-affinity binders (T4 and T7) were identified and tested for complex formation using *Fn*SiaQM. Hexa-histidine-tag based affinity chromatography was used to purify the nanobody from the host bacterial periplasm, producing a monodisperse peak in the size-exclusion chromatogram. Both nanobodies were added to *Fn*SiaQM in DDM micelles and tested for complex formation. Although both nanobodies bound to the transporter, the T4 nanobody was selected over T7 because of its higher expression. The structures of the purified *Fn*SiaQM-nanobody (T4) complexes in nanodiscs with and without Neu5Ac bound were determined using cryo-EM.

### Structure of the Fn*SiaQM*-nanobody complex

The nanodisc reconstituted *Fn*SiaQM, and the bound nanobody is a monomeric complex in the cryo-EM structure. The overall global resolutions of the both the unliganded *Fn*SiaQM-nanobody complex and the Neu5Ac bound form is ∼3.2 Å. The maps were of good quality and allowed us to build all the TM helices of FnSiaQM unambiguously and both the unliganded and the Neu5Ac bound form clearly allow us to trace the protein and the nanobody **(Figure 1A**, **1B**, **1C**, **1D**). The best resolution for both structures are observed in the interior of the SiaQM protein (**Supplementary Figure 3C, 3D**). The details of the structure quality and refinement parameters are shown in **Table 1**, and a flow diagram for structure determination and resolution statistics is shown in **Supplementary Figure 3 A and B**. Owing to their flexibility, the N-terminal 6X histidine and Strep II tags on *Fn*SiaQM-nanobody complex were not visible in the cryo-EM maps. Similarly, the last few amino acids at the C-terminus were not built because of poor density. The density of the nanodisc was visible but of insufficient quality for model building. The density of the bound Neu5Ac is clear and in the higher-resolution region. The observed density around the ligand binding site for the two maps clearly indicate minimal structural changes around the binding pocket (**Figure 1C**, **1D**, inset).

**Figure 1:**
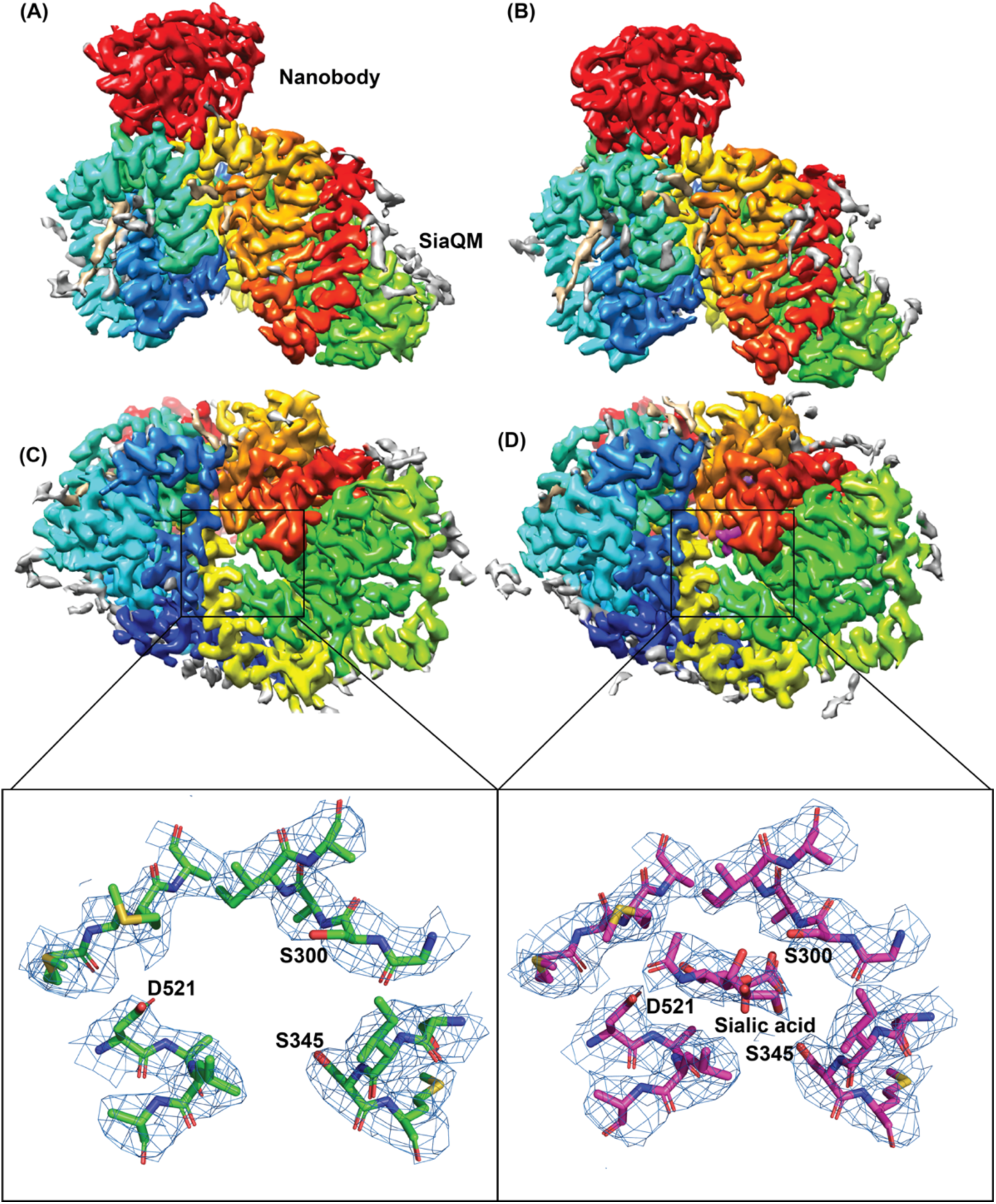
Architecture of FnSiaQM with nanobody. (**A and B**) Cryo-EM maps of FnSiaQM unliganded and Neu5Ac bound at 3.2 and 3.17 Å, respectively. The TM domain of FnSiaQM is colored using the rainbow model (N-terminus in blue and C-terminus in red). The nanobody density is colored in red. The density for modeled lipids is colored in tan and the unmodelled density in gray. The figures were made with Chimera at thresholds of 1.2 and 1.3 for the unliganded and Neu5Acbound maps. (**C and D**) The cytoplasmic view of apo and Neu5Ac bound FnSiaQM, respectively. Color coding is the same as in panels A and B. The density corresponding to Neu5Ac and sodium ions are in purple. The substrate binding sites of apo and Neu5Ac bound FnSiaQM are shown with key residues labeled. The density (blue mesh) around these atoms was made in Pymol with 2 and 1.5 σ for the apo and the Neu5Ac structures, respectively, with a carve radius of 2 Å.

As the name suggests, many TRAP transporters comprise three units: a substrate-binding protein (SiaP) and two membrane-embedded transporter units (SiaQ and SiaM) (Severi *et al*., 2007). In *Fn*SiaQM, the two transporter units are fused by a long connecting helix, similar to other TRAP transporters such as *H. influenzae* TRAP (*Hi*SiaQM) (Currie *et al*., 2024; Peter *et al*., 2022). The Q-domain of *Fn*SiaQM consists of the first four long helices, which are tilted in the membrane plane. This tilting creates a large contact surface area between the Q- and M-domains, which is dominated by buried hydrophobic residues. A small connecting helix lies perpendicular to the cell membrane plane and does not contact either domain (**Figure 2**). The lack of interaction between this connecting helix and the two domains suggests that it may be redundant for the transporter function. For example, *Pp*SiaQM contains two separate polypeptides and does not require a connecting helix for its function (Davies *et al*., 2023).

**Figure 2:**
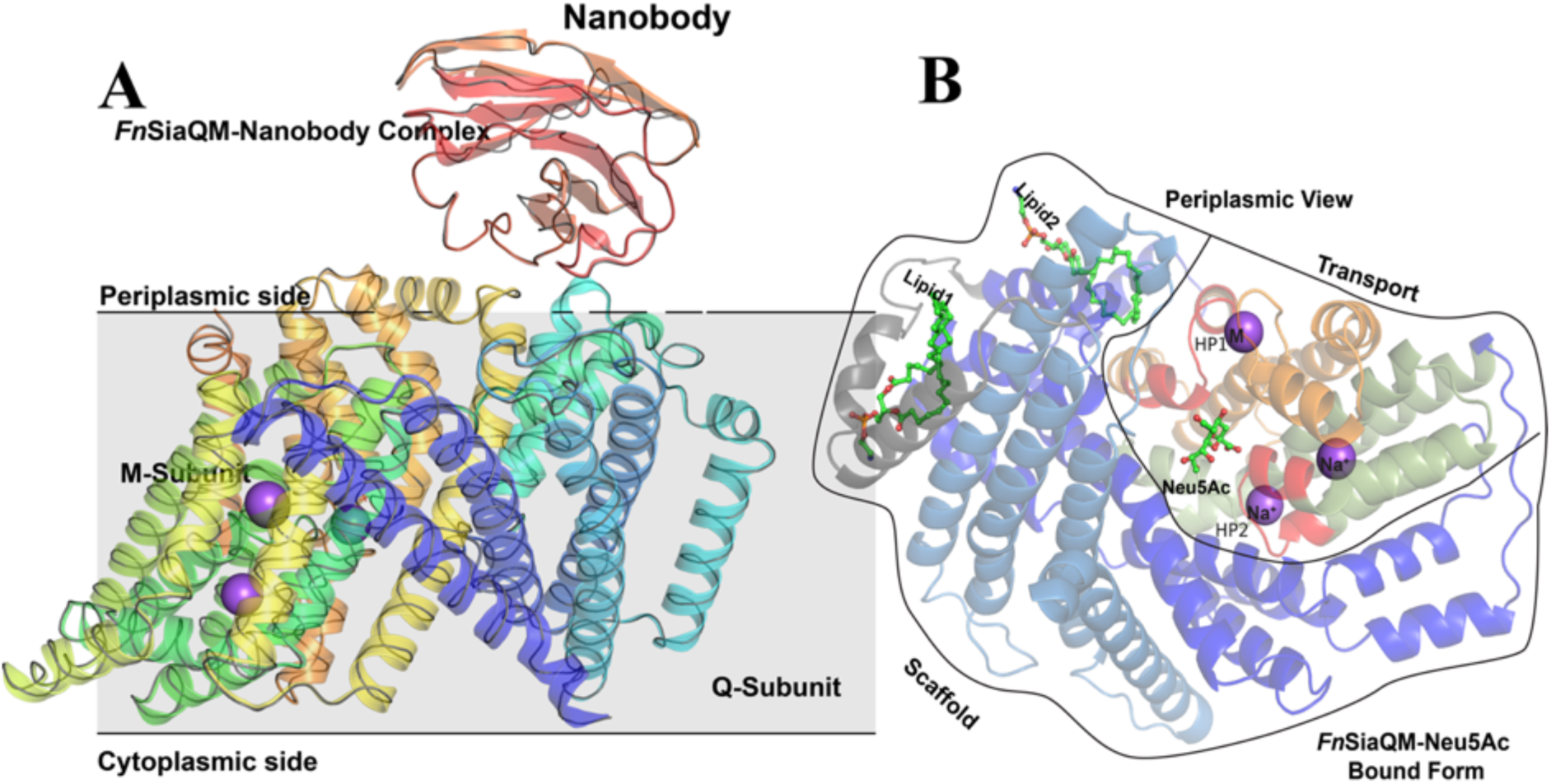
*Fn*SiaQM-Nanobody complex with Neu5Ac bound form. (**A**) A cartoon representation of the *Fn*SiaQM-Nanobody complex structure. The SiaQM polypeptide has been colored in rainbow with the N-terminus starting in blue and the C-terminus ends in orange. The nanobody is shown in cartoon representation and in red color. Two of the modeled sodium sites are shown as purple spheres. A ribbon representation of the unliganded structure of *FnSiaQM* is superposed in grey. The superposition reveals that the overall structures are similar. (**B**) Cartoon representation of the *Fn*SiaQM structure bound to Neu5Ac. In shades of blue are the helices that form the scaffold domain. In gray is the connecting domain. In olive, orange, and red is the elevator domain. HP1 and HP2, the two helix-loop-helix motifs, are in red and marked. The positions of the known Na^+^ ions and metal ion (M) are in spheres, and the position of Neu5Ac is shown in ball and stick.

While there is no direct observation, the possible mode of function of the transporter suggests that the N-terminus of *Fn*SiaQM lies on the cytoplasmic side, whereas its C-terminus lies on the periplasm side of the membrane. The nanobody is bound to the periplasmic side of *Fn*SiaQM, making no contact with the cytoplasmic side of the protein (**Figure 1A and 1B**). The M-domain is located towards the C-terminus and comprises the bulk of the transporter. It consists of 10 transmembrane helices with two hairpins that do not cross the entire membrane.

Superposition of the reported structures from *Hi*SiaQM and *Pp*SiaQM demonstrate that all are in inward (cytoplasmic) open and elevator-down conformations (**Supplementary Figure 1**). There are two conserved Na^+^ ion binding sites in these proteins (Currie *et al*., 2024; Davies et al., 2023; Peter *et al*., 2022). Structurally, the M-domain can be divided into an outer variable scaffold domain and a centrally conserved transport domain. The scaffold domain is composed of six TM helices and serves as a support for the transport domain. It also interacts with the Q-domain to form a structurally rigid domain. The Q and scaffold domains are the least conserved among the SiaQM domains, suggesting that they are not directly involved in substrate transport (**Figure 2** and **Supplementary Figure 4**).

To determine whether substrate binding resulted in conformational changes, we solved the structure of the ligand-bound form of *Fn*SiaQM. The superposition of the two structures revealed a high degree of similarity with an RMSD of 0.3 Å over all Cα atoms (**Figure 2A**). A comparison of the binding site residues revealed that they are in a similar conformation. The transporter has a large cytoplasmic facing open cavity, suggesting that the unliganded as well as Neu5Ac bound structures are in an inward-open conformations. The nomenclature used for labeling the secondary structures uses a description of the published *Hi*SiaQM structure with similar architecture (Currie *et al*., 2024). The elevator part of the domain has one helix-loop-helix motif called HP1 (HPin), which is directed towards the cytoplasmic side. A structurally homologous helix-loop-helix domain, HP2 (HPout), is present on the periplasmic side. The arrangement of TM helices have been described in detail in *Hi*SiaQM and *Pp*SiaQM structures (Davies *et al*., 2023; Peter *et al*., 2022). Interestingly, this domain has two-fold inverted symmetry, as found in other TRAP transporters. The entire domain is populated with highly conserved residues, suggesting that it may play a direct role in substrate transport (**Supplementary Figure 4**). A single molecule of Neu5Ac is bound to the aperture formed by HP1 and HP2 in the central core transport domain (**Figure 2B**). The Neu5Ac binding site has a large solvent-exposed vestibule towards the cytoplasmic side, while its periplasmic side is sealed off. Cryo-EM map shows the presence of multiple densities that could be modeled as lipids, possibly preventing the substrate from leaving the transporter. However, the densities are not well defined to model them as specific lipids, hence they have not been modeled. We describe this as the “inward-facing open state” with the substrate-bound. While, both the HP1 and HP2 loops have been hypothesized to be involved in gating, in the human neutral amino acid transporter (ASCT2), (which also uses the elevator mechanism), only the HP2 loops have been shown to undergo conformational changes to enable substrate binding and release(Garaeva *et al*., 2019). Hence, it is suggested that there is a single gate that controls substrate binding. Superposition of the *Pp*SiaQM and *Hi*SiaQM structures do not reveal any change in these loop structures upon substrate binding. For TRAP transporters, the substrate is delivered to the QM protein by the P protein; hence, these loop changes may not play a role in ligand binding or release. This may support the idea that there is minimal substrate specificity within SiaQM and that it will transport the cargo delivered by SiaP, which is more selective.

### Sodium ion binding site

The transport of molecules across the membrane by TRAP transporters depends on Na^+^ transport (Currie *et al*., 2024; Davies *et al*., 2023; Mulligan *et al*., 2012, 2009). We observed cryo-EM density at conserved Na^+^ binding sites and modeled two Na^+^ ions in the unliganded- and ligand-bound forms (**Figure 3**). The two sites that are present in the unliganded form are also conserved in other reported SiaQM structures (Currie *et al*., 2024; Davies *et al*., 2023) (**Figure 2B, Supplementary Figure 5)**, compared with Figure 5 in (Currie *et al*., 2024)]. The Na1 site interacts with residues at the HP1 site and helix 5 (**Figure 3A, Supplementary Figure 6A**). The Na2 site combines residues at the HP2 site and a short stretch that splits helix 11 into two parts (**Figure 3A, Supplementary Figure 6B**). These two sites have been previously well described (Currie *et al*., 2024). These two sodium ion binding sites are also conserved in the structure of VcINDY (**Supplementary Figure 7)** (Sauer *et al*., 2022). In both cases, the sodium ions are bound at the helix-loop-helix ends of HP1 and HP2. The binding sites utilize both side chains and main chain carbonyl groups. The number of main chain carbonyl interactions suggests that they are critical, and using main chain rather than side chain interactions minimizes the likelihood of point mutations affecting the binding.

**Figure 3:**
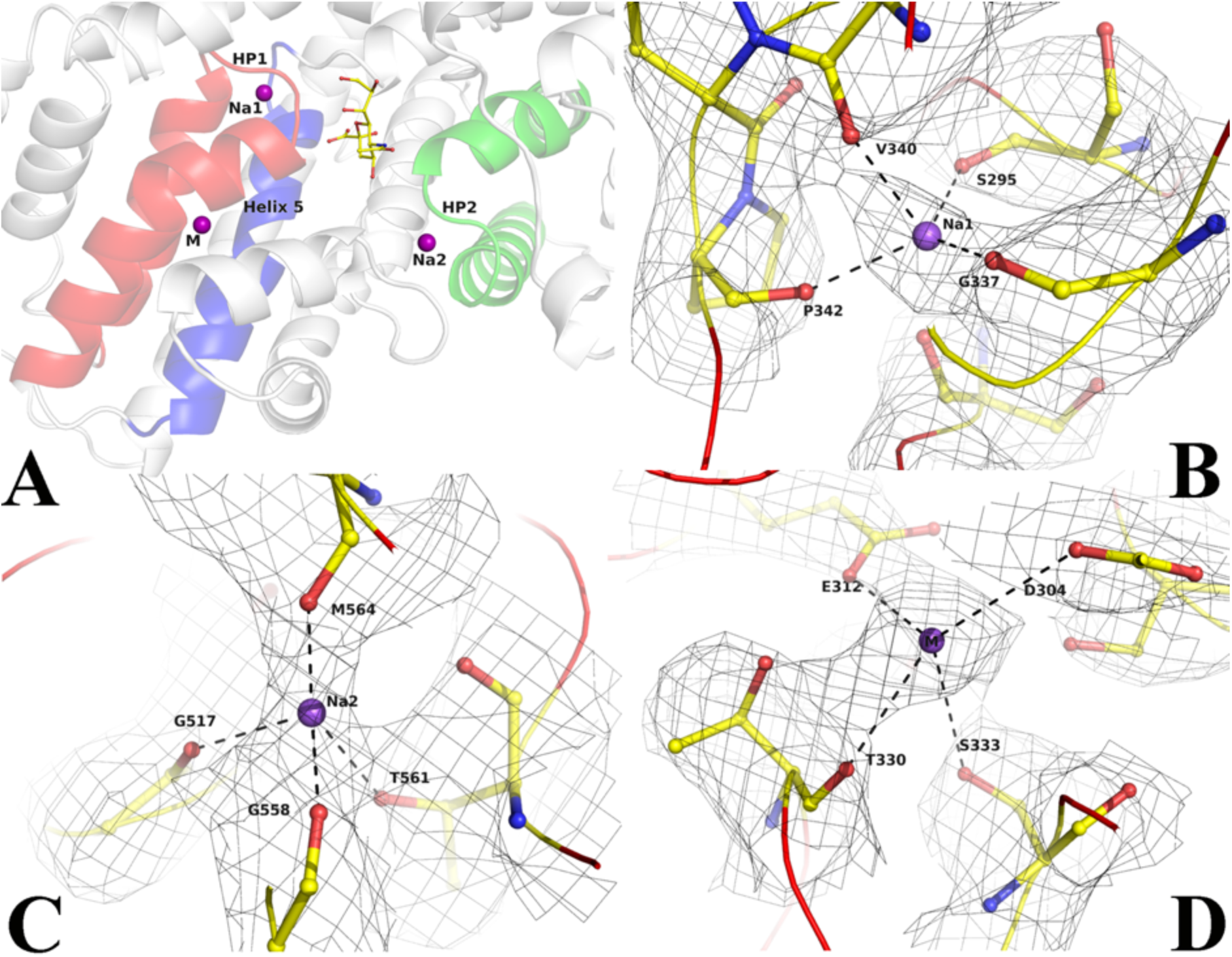
Ion binding sites in *Fn*SiaQM. (**A**) Close-up view of the Neu5Ac and the two Na^+^ and metal binding sites. In red is the HP1 helix-loop-helix. In green is the HP2 helixloop-helix. In blue is helix 5 A, the purple spheres are the two Na^+^ ion binding sites and metal binding site (M), and the bound Neu5Ac is shown in ball and stick. (**B-D**) Density and interaction details of the Na1, Na2, and M sites, respectively. The figures are made with a contour of 1.0 r.m.s. in PyMol.

Surprisingly, we observed density for an additional metal binding site. This site is distinctly located towards the cytoplasmic side of the transporter and away from the Neu5Ac binding site. It interacts with helices that form HP1 and the next helix 5a (**Figure 3A**). Interestingly, a similar site was observed in the SSS Neu5Ac transporter, in which the third site was located away from the other Na^+^ binding sites (Wahlgren *et al*., 2018). Peter *et al.,* proposed that Asp304 coordinates with this Na^+^ ion, showing loss-of-function when mutated to an alanine residue (Peter *et al*., 2022). While it is tempting to label it as a third Na^+^-binding site, the metal– ligand distances are longer (**Supplementary Figure 6C**). Hence, we conservatively designated this site as a metal-binding site. The coordinated movement of different Na^+^ ions is expected to provide the energy to create conformational changes that lead to the movement of Neu5Ac from the periplasm to the cytoplasm. While we found two Na^+^-binding sites and a third metal-binding site, it is not clear how the movement of these sites can cause the conformational changes required for the motion of the elevator along with the ligand. It is also unclear what the function of the third metal-binding site is beyond the mutagenesis data, which suggests that the lack of this site results in a loss of function.

### Lipid binding to *Fn*SiaQM

We have modelled two phosphatidylethanolamine (PE) lipids in the maps of unliganded *Fn*SiaQM (**Supplementary Figure 8A and 8B**). The *Fn*SiaQM nanodisc reconstitution procedure included the addition of *E. coli* polar lipids, where PE is the most abundant lipid. One of the lipids binds between the connector helix and the Q-domain (**Figure 2B, Lipid1**) and is present in both the unliganded and Neu5Ac bound structure of *Fn*SiaQM. This lipid is present at a similar position in the *Hi*SiaQM structure (Currie *et al*., 2024). However, we observe a second lipid molecule (**Figure 2B, Lipid2**) that is bound between the stator and elevator domains only in the Neu5Ac bound structure. It is tempting to hypothesize that this lipid molecule is displaced during the movement of the elevator domain; however, this requires further investigation. A similar observation was also made in the transporter GltPh that invokes the elevator mechanism (Wang and Boudker, 2020).

### Activity of *Fn*SiaQM

To demonstrate that the purified protein is active and transports Neu5Ac across the membrane, we performed transport assays. *Fn*SiaQM was reconstituted in proteoliposomes containing intraliposomal potassium. Valinomycin was then incorporated into these proteoliposomes to establish an inner-negative membrane potential. Periplasmic binding protein (*Fn*SiaP) and radioactive Neu5Ac were added to the proteoliposomes to initiate transport (**Figure 4A**). In this experimental setup, a significant accumulation of radiolabeled Neu5Ac was measured in the presence of extraliposomal Na^+^-gluconate when the artificial membrane potential was imposed by the addition of valinomycin (**Figure 4B, black squares**). When ethanol was used as a control for valinomycin, a much lower accumulation of radiolabeled Neu5Ac was observed (**Figure 4B, open squares**). Importantly, transport was negligible when Na^+^ was absent in the extra-liposomal environment or when K^+^ was absent in the intraliposomal environment. Transport was negligible when soluble *Fn*SiaP was omitted from the assay, further demonstrating the requirement of periplasmic binding proteins (**Figure 4B, stars**). The energetics of transport is similar to that of other SiaPQM that have been characterized (Mulligan *et al*., 2012).

**Figure 4:**
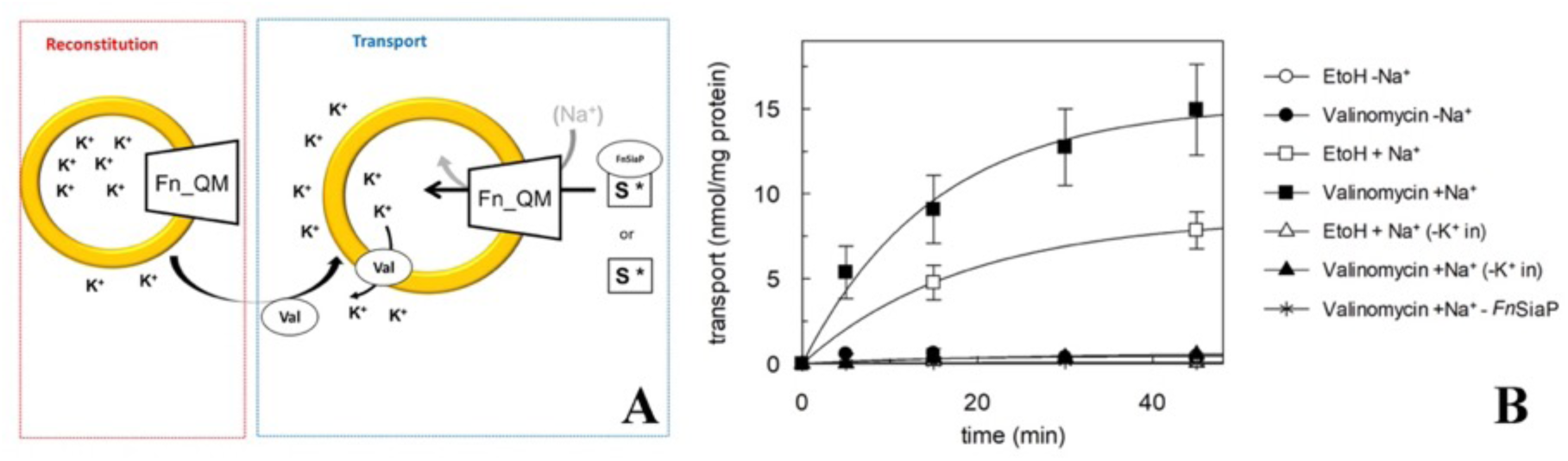
Proteoliposome transporter assays for *Fn*SiaQM. (**A**) Schematic diagram showing the experimental setup. First, *Fn*SiaQM is incorporated into proteoliposome in the presence of internal K^+^. Then, valinomycin (val) is added to induce the efflux of K^+^ down its concentration gradient, imposing an artificial membrane potential. To start transport measurement, *Fn*SiaP, [^3^H]-Neu5Ac and Na^+^-gluconate are added in the extraliposomal environment. (**B**) Time course of Neu5Ac uptake into proteoliposomes reconstituted with *Fn*SiaQM. In black circle, black square, white triangles, black triangle, asterisk, are with conditions, where valinomycin was added to facilitate K^+^ movement before transport. Ethanol was added instead of valinomycin as a control in the white circle, white square, and white triangle. In white square, black square, white triangle, black triangle, 10 mM Na^+^-gluconate was added together with 5 µM [^3^H]-Neu5Ac and 0.5 µM *Fn*SiaP; in white triangle and black triangle proteoliposomes are prepared without internal K^+^; in black asterisk transport is measured in the presence of 10 mM Na^+^-gluconate, 5 µM [^3^H]-Neu5Ac and in the absence of *Fn*SiaP. Uptake data were fitted in a first-order rate equation for time course plots. Data are means ± s.d. of three independent experiments.

### Architecture of Neu5Ac binding site

While the structures of the unliganded and Neu5Ac bound *Fn*SiaQM are in similar inward-open conformation, the density of Neu5Ac is unambiguous in the liganded form (**Figure 1D (inset), 5A**). The overall architecture of the Neu5Ac binding site is similar to that of citrate/malate/fumarate in the di/tricarboxylate transporter of *V. cholerae* (*Vc*INDY), but the residues involved in providing specificity are different (Kinz-Thompson *et al*., 2022; Mancusso *et al*., 2012; Nie *et al*., 2017; Sauer *et al*., 2022). Neu5Ac binds to the transport domain without direct interactions with the residues in the scaffold domain. The majority of the interactions are with residues in the HP1 and HP2 loops of the transport domain (**Figure 5B**). Asp521 (HP2), Ser300 (HP1), and Ser345 (helix 5) interact with the substrate through their side chains, except for one interaction between the main chain amino group of residue 301 and the C1-carboxylate oxygen of Neu5Ac. Mutation of the residue equivalent to Asp521 has been shown to result in loss of transport (Peter *et al*., 2022). To evaluate the role of residues Ser-300 and Ser-345, we mutated them to alanine and performed the transport assays.

**Figure 5:**
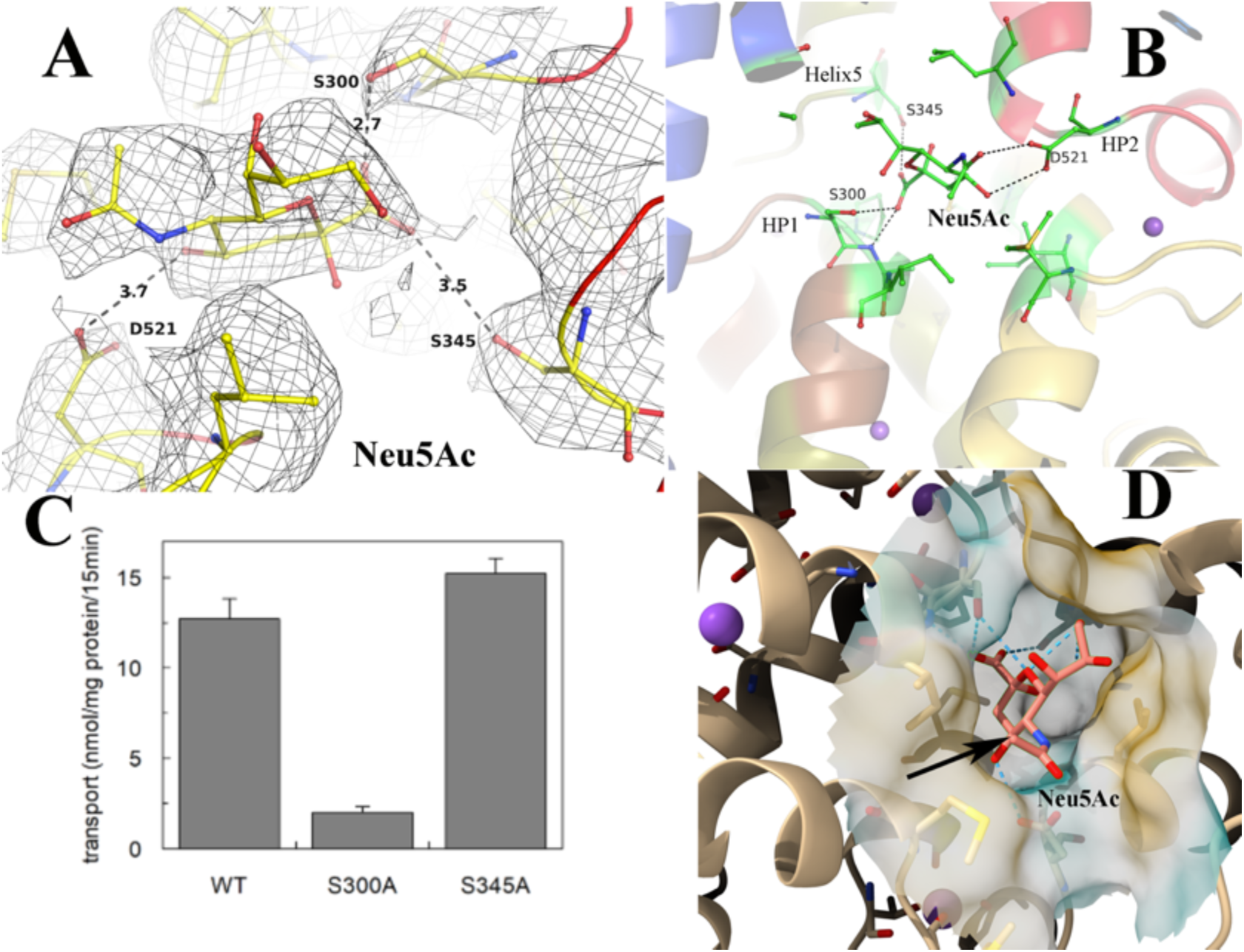
Validation of Neu5Ac binding pocket in *Fn*SiaQM transporter. (**A**) The fit of the modeled Neu5Ac into the density, contoured to 0.9 * r.m.s. The figure also shows the fit to the density of the residues that interact with Neu5Ac and the distances to key residues discussed in the manuscript. (**B**) The interactions of Neu5Ac with side chains of interacting residues. Ser300 Oγ is 2.8 Å from the C1-carboxylate oxygen of Neu5Ac, while the main chain NH of residue 301 is 2.6 Å away. The other close polar side chain is that of Ser345γ, which is 3.3 Å away. The Neu5Ac O10 is 2.9 Å from Asp521 Oδ. (**C**) Transport of Neu5Ac in 15 min. Transport was started by adding 5 µM [^3^H]-Neu5Ac, 10 mM Na^+^-gluconate, and 0.5 µM *Fn*SiaP to proteoliposomes in the presence of valinomycin (expected membrane potential, ΔΨ, −117.1 mV). Data are expressed as nmol/mg prot/15 min ± s.d. of three independent experiments. (**D**) The cavity of Neu5Ac is exposed from the cytoplasmic side. The cavity is large and extends into the cytoplasmic side. The black arrow shows the methyl group with an extra hydroxyl in Neu5Gc.

The data clearly showed that the Ser300Ala mutant was inactive, whereas the Ser345 mutation did not affect *Fn*SiaQM functionality (**Figure 5C**). This suggests that the interaction of Neu5Ac with the residues in HP1 and HP2 is critical for transport. The carboxylate oxygens of the C1 atom of Neu5Ac interact with Ser300 and Ser345 and the main chain of Ala301 (**Figure 5B**). Ser 345 OG is 3.5Å away from the C1-carboxylate oxygen – a distance that would result in a weak interaction between the two groups. It is, therefore, not surprising that the mutation into Ala did not affect transport. The space created by the mutation can be occupied by a water molecule. The N5 atom of the *N*-acetyl group of the sugar interacts with Asp521. These interactions are conserved even if Neu5Ac is converted into Neu5Gc, as the addition of an extra hydroxyl group at the C11 position does not break this interaction, and there is sufficient cavity space for modifications in C11 (**Figure 5D**). Cavity Plus (Wang *et al*., 2023) estimated the cavity to be 1875 Å^3^ with a druggability score of 3631, suggesting that the environment of the binding site is highly suited for drug binding (Wang *et al*., 2023).

No direct measurement of *K*_d_ of Neu5Ac binding to the SiaQM region of the transporter is available. We infer from the interactions it makes and the fact that high concentrations of Neu5Ac are required to obtain a complex (30 mM), the affinity is unlikely to be high. Neu5Ac binds with nanomolar affinity (K_d_ = 45 nM) to *Fn*SiaP (**Figure 6A**) and several electrostatic interactions stabilize this binding (Gangi Setty *et al*., 2014). The binding of Neu5Ac to the SSS-type transporter, which also transports Neu5Ac across the cytoplasmic membrane, was also stabilized by several electrostatic interactions (**Figure 6B**). The polar groups bind to both the C1-caboxylate side of the molecule and the C8-C9 carbonyls, suggesting that *Proteus mirabilis* Neu5Ac transporter (SSS type) evolved specifically to transport nine-carbon sugars such as Neu5Ac (Wahlgren *et al*., 2018). Interestingly, even the dicarboxylate transporter from *V. cholerae* (*Vc*INDY) binds to its ligand via electrostatic interactions with both carboxylate groups (**Figure 6C**) (Kinz-Thompson *et al*., 2022; Mancusso *et al*., 2012; Nie *et al*., 2017; Sauer *et al*., 2022). The high affinity of the substrate-binding component (*Fn*SiaP) to Neu5Ac is physiologically relevant because it sequesters Neu5Ac in a volume where the concentration of Neu5Ac is very low. In contrast, as Neu5Ac is delivered to the SiaQM component by the SiaP component, the affinity could be lower, an argument also made by Peter *et. al*. in their recent work (Peter *et al*., 2024). The corollary of this argument is that the specificity of the transport system (or the choice of molecules that can be transported) is likely to be determined by the substrate-binding component. There is probably very little selectivity in the SiaQM component, which is also reflected by fewer interactions.

**Figure 6:**
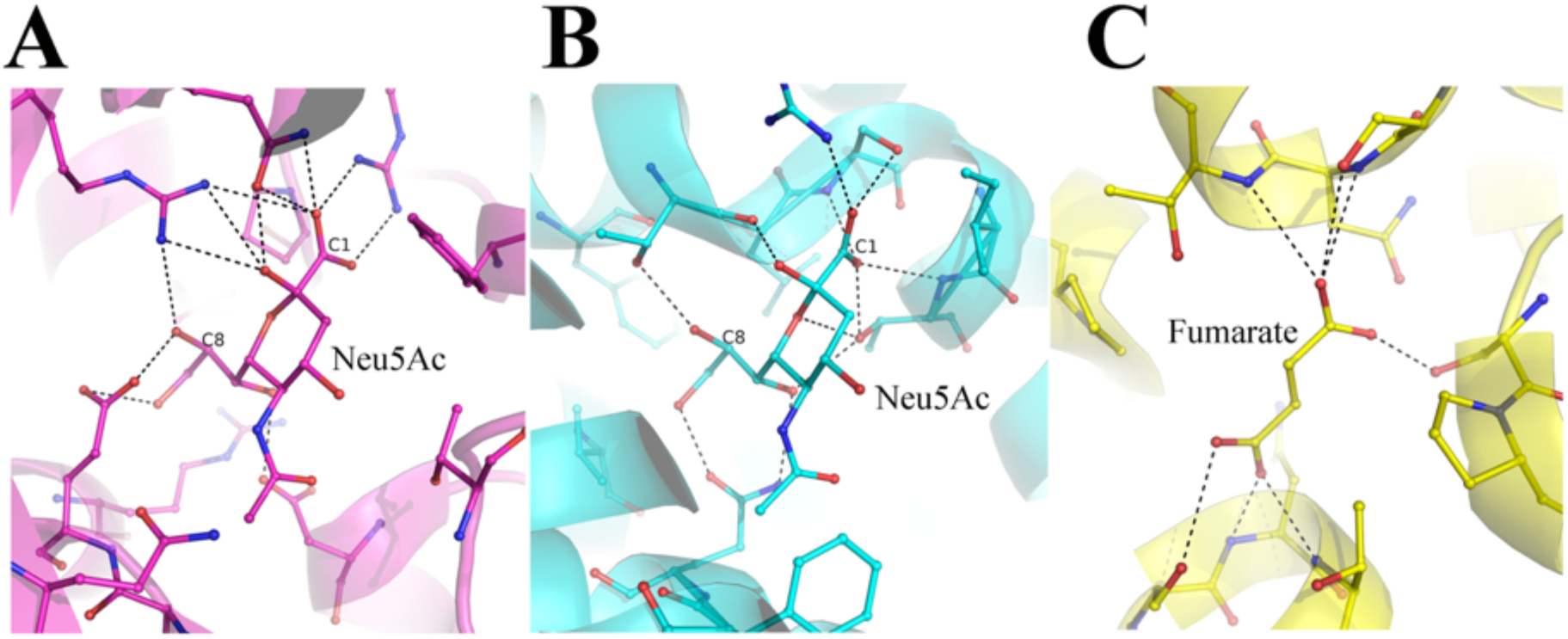
Comparison of Neu5Ac binding pocket. (**A**) The interactions of Neu5Ac with the substrate binding protein SiaP from *F. nucleatum* (PDB-ID 4MNP). The C1 and the C8 carbon atoms are labeled to show the different ends of the nine-carbon sugar. (**B**) the interaction of Neu5Ac with the SSS-type Neu5Ac transporter (PDB-ID 5NV9). (**C**) The interaction of fumarate with the dicarboxylate transporter *Vc*INDY (PDB-ID 6OKZ).

## Conclusion

This is the first report of the structure of a SiaQM from the TRAP-type transporter of Gram-negative bacteria with bound Neu5Ac. The structure shows no direct interaction between Neu5Ac and Na^+^ ions. The affinity of Neu5Ac for periplasmic sialic acid-binding proteins (SiaP) is in the nanomolar range, with many polar interactions (K_d_=45 nM) (Gangi Setty *et al*., 2014). However, relatively fewer polar interactions stabilize Neu5Ac binding to the open binding pocket of SiaQM, and the affinity is probably poor. We used APBS to calculate the electrostatic charge distribution on SiaQM **(Supplementary Figure 8C)** (Jurrus *et al*., 2018). A view of the structure of Neu5Ac bound SiaQM from the periplasmic side towards the bound Neu5Ac shows that the sugar is bound in the cavity, the entrance of which is repulsive to the binding of a negatively charged sugar like Neu5Ac. This likely prevents Neu5Ac from binding to the SiaQM from the cytoplasmic side. The structure of the protein in the outward-facing conformation is unknown, which will reveal how Neu5Ac is transferred from SiaP to SiaQM. While mechanistic questions require further investigation, the precise definition of the binding pocket provided in this work is a starting point for structure-based drug discovery.

## Experimental procedures

### Construct design

The gene sequence encoding the TRAP transporter from *F. nucleatum* (NCBI Reference Sequence: WP_005902322.1) was synthesized and cloned into a pBAD vector by GeneArt. A Strep-tag was inserted in addition to the 6X-His Tag at the N-terminus to enhance the protein purity. It was also cloned into pWarf (+) vector for fluorescence-detection size-exclusion chromatography (Hsieh *et al*., 2010).

### Protein purification of *Fn*SiaQM

Expression trials and detergent scouting were performed as described by Hsieh *et al*., 2010. For large-scale production, the *Fn*SiaQM construct in pBAD was introduced into *E. coli* strain TOP10 cells. A single colony from the transformed plate was inoculated into 150 mL of terrific broth (TB) containing 100 µg/mL ampicillin. The culture was then incubated overnight at 30 °C in a shaker incubator. The following day, 10 mL of the primary culture was transferred to a culture flask containing 1 L Terrific broth supplemented with ampicillin and incubated at 30 °C in an orbital shaker. The cultures were induced with 0.02% arabinose at an optical density (OD) of 1.5 at 600 nm and allowed to grow for an additional 18 h. Cells were harvested by centrifugation at 4000 x g for 20 min at 4 °C. The resulting cell pellet was rapidly frozen in liquid nitrogen and stored at −80 °C for future use. The cell pellet was resuspended in lysis buffer (100 mM Tris, pH 8.0, 150 mM NaCl, and 1 mM dithiothreitol (DTT)) supplemented with lysozyme, DNase, and cOmplete EDTA-free inhibitor cocktail (Roche). Lysis was achieved by two passes through a Constant cell disruptor at 20 and 28 kPsi, respectively. Subsequently, cell debris and unbroken cells were pelleted by centrifugation at 10,000 x g for 30 min at 4 °C. The supernatant from the low-speed centrifugation was subjected to ultracentrifugation at 150,000 x g at 4 °C for 1 h to pellet the cell membranes. The membranes were then resuspended in solubilizing buffer (1% n-dodecyl-β-D-maltoside (DDM), 50 mM Tris pH 8.0, 150 mM NaCl, 10 mM imidazole, 3 mM β-ME) at a 1:10 w/v ratio and solubilized at 4 °C for two hours. The insoluble membrane components were separated by ultracentrifugation for 30 min at 100,000 × g, and the resulting supernatant was loaded onto a 5 mL His-Trap (Cytiva) column pre-equilibrated with buffer A (50 mM Tris pH 8.0, 150 mM NaCl, 10 mM imidazole, 3 mM β-ME, 0.02% DDM). The resin was extensively washed with buffer A until a stable UV absorbance was achieved in the chromatogram. The *Fn*SiaQM protein was eluted using an elution buffer (50 mM Tris, pH 8.0, 150 mM NaCl, 150 mM imidazole, 3 mM β-ME, and 0.02% DDM). The eluted fractions were pooled and used for strep-tag purification. A 5 mL Strep-Trap HP column (Cytiva) was used for subsequent purification. The column was pre-equilibrated with a buffer (100 mM Tris, pH 8.0, 150 mM NaCl, 10 mM imidazole, 3 mM β-ME, and 0.02% DDM). After loading the sample onto the column, the column was washed with 10 CV buffer (100 mM Tris pH 8.0, 150 mM NaCl, 10 mM imidazole, 3 mM β-ME, and 0.02% DDM). The *Fn*SiaQM protein was eluted in the presence of 10 mM desthiobiotin, 100 mM Tris (pH 8.0), 150 mM NaCl, 3 mM β-ME, and 0.02% DDM. The eluted fraction was concentrated using a 100 kDa Amicon filter and subsequently injected into a size-exclusion column (Superdex200 16/300 increase) in buffer (50 mM Tris pH 8.0, 150 mM NaCl, 1 mM DTT, 0.02% DDM). Size exclusion chromatography yielded a monodisperse peak for *Fn*SiaQM protein (**Supplementary Figure 2**).

### Generation, isolation and purification of *Fn*SiaQM-specific nanobodies

Generation of the nanobodies was performed by the University of Kentucky Protein Core, following previously established protocols (Chow *et al*., 2019). Alpacas were subcutaneously injected with 100 μg of recombinant *Fn*SiaQM in DDM once a week for six weeks. Peripheral blood lymphocytes were isolated from alpaca blood and used to construct a bacteriophage display cDNA library. Two rounds of phage display against the recombinant *Fn*SiaQM identified two potentially VHH-positive clones. Positive clones were confirmed by sequencing and were analyzed for nanobody components. These two nanobodies were cloned into the pMES4 vector and *E. coli* strain BL21 (DE3) was used for their expression. A single colony was inoculated into 100 mL LB medium and cultured overnight. These overnight cultures were diluted in 1 L of LB media in a 1:100 ratio. Expression was induced with 0.5 mM Isopropyl β-D-1-thiogalactopyranoside (IPTG) when the O.D._600_ reached 1.0, followed by further incubation at 28 °C for 16 h for protein production. Cultures were then centrifuged at 4000 x g for 20 min at 4 °C, and periplasmic extracts were prepared using the osmotic shock method with 20% sucrose. The periplasmic extract was dialyzed to remove sucrose and filtered for affinity chromatography. This filtered fraction was loaded onto a 5 mL His-Trap (Cytiva) column pre-equilibrated with buffer A (50 mM Tris pH 8.0, 150 mM NaCl, 10 mM imidazole, and 3 mM β-ME). The column was extensively washed with 20 CV of buffer B until stable UV absorbance was observed in the chromatogram. The nanobodies were eluted using an elution buffer (50 mM Tris, pH 8.0, 150 mM NaCl, 150 mM imidazole, and 3 mM β-ME). The eluted fractions were pooled and concentrated using a 10 kDa Amicon (Sigma) filter and subsequently injected into a Superdex75 column. Both nanobodies exhibited monodisperse peaks during the size-exclusion chromatography. The peak fractions were pooled and stored at 4 °C.

### Complex formation of *Fn*SiaQM-nanobody (T4)-nanodisc (MSP1D1)

MSP1D1 was expressed and purified following established protocols (Denisov and Sligar, 2016). The 6X-His tag was cleaved from the affinity-purified MSP1D1 using TEV protease and the resulting product was concentrated to 5 mg/mL. For nanodisc formation, nickel affinity-eluted *Fn*SiaQM, MSP1D1, and *E. coli* polar lipids were combined in a molar ratio of 1:6:180 and incubated on ice for 15 min. Nanodisc formation was initiated by adding washed Bio-Beads (200 mg dry weight per 1 mL of the protein mixture; Bio-Rad), and the mixture was rotated for two hours at 4 °C. The protein mixture was subsequently separated from the Bio-Beads using a fine needle. A two-molar excess of the T4-Nanobody was added at this step to the *Fn*SiaQM-nanodiscs. A Strep-Tag chromatography step removed the empty nanodiscs and excess T4-nanobodies. The *Fn*SiaQM-reconstituted nanodisc with the T4 nanobody were then separated from the aggregates using a Superdex200 16/300 increase size exclusion column. The fractions corresponding to the peaks were concentrated to 2.5 mg/mL and used for cryo-EM analysis. To acquire the Neu5Ac-bound structure, 30 mM Neu5Ac was incorporated in the gel filtration buffer (100 mM Tris, pH 8.0, 150 mM NaCl, and 1 mM DTT).

### Cryo-EM sample preparation and data collection

The *Fn*SiaQM-T4 complex at a concentration of 2.5 mg/mL was immediately applied onto UltrAuFoil R0.6/1.0 grids (300 mesh) previously subjected to glow discharge for 150 s at 25 mA. Excess fluid was removed by blotting using a Vitrobot Mark IV (ThermoFisher Scientific) apparatus, and the grids were rapidly vitrified by plunging them into liquid ethane that was pre-cooled with liquid nitrogen. Subsequently, the grids were screened at a magnification of 59,000 × using a Titan Krios microscope G3i (ThermoFishser Scientific) operating at 300 kV. Data were collected at a magnification of 75,000 ×, corresponding to a pixel size of 1.07 Å using a Falcon 3 detector in counting mode.

### Data processing of unliganded form

A total of 960 movies were collected for the unliganded structure. All processing was performed using the CryoSPARC software suite (Punjani *et al*., 2017). After patch motion correction and CTF correction, the summed images were examined and 941 micrographs were selected for particle picking. The initial particle picking (blob picker) was performed using 315 images. This was followed by 2D classification, the selection of 2D classes, and *ab initio* reconstruction into two classes. One class clearly showed the presence of the nanodisc, the protein inside, and the bound nanobody. This was refined with 49,659 particles to a resolution of 6.7 Å resolution. This 3-D map was used to create templates, template-based particle picking was performed on all 941 images, and 385,668 particles were picked. Multiple rounds of 2-D classification and selection yielded 141,272 good particles. These particles were used for *ab initio* reconstruction in cryoSPARC (Punjani *et al*., 2017), followed by homogenous and non-uniform refinement (Punjani, 2020). The final map resulted in an overall resolution of 3.2 Å resolution (Fourier Shell Correlation cutoff 0.143). The B-factor estimated from the Guinier plot is 125.2 Å^2^.

### Data processing of Neu5Ac-bound form

A total of 2341 images were collected. After manually inspecting the micrographs, 1752 images were selected for further processing. The template generated from the unliganded maps was used for the particle picking. After multiple rounds of 2D classification and particle pruning, 653,554 particles were used to create six *ab initio* classes. Four of these classes appeared identical (225,006 particles) and were refined to better than 4.5 Å using homogeneous refinement. The final map was constructed using non-uniform refinement protocols in Cryo-SPARC. The map has an overall resolution of 3.17 Å at the FSC 0.143 threshold. The B-factor estimated from the Guinier plot is 135.5 Å^2^.

### Model building and model refinement

The starting model was constructed using Alphafold2 software (Evans *et al*., 2022). Subsequently, this model was docked within the cryo-EM map of *Fn*SiaQM unliganded form of *Fn*SiaQM. A series of iterative rounds of model building using Coot (Emsley and Cowtan, 2004) and refinement using Phenix (Adams *et al*., 2010) were performed to complete the model. These steps were essential for enhancing the accuracy and quality of the structural model and ensuring its congruence with the experimental cryo-EM data. The refined unliganded model was used as the starting model for the substrate-bound map, and sugars and ions were modeled. All the figures were either made in UCSF Chimera or PyMOL (DeLano, 2002; Pettersen *et al*., 2004). The final details of the data collection, processing, and results of model building and refinement are summarized in **Table 1**.

### Reconstitution of *Fn*SiaQM into proteoliposomes

Purified *Fn*SiaQM was reconstituted using a detergent removal method performed in batches following previously described procedures (Wahlgren *et al*., 2018). In summary, 50 μg *Fn*SiaQM was combined with 120 μL of 10% C_12_E_8_ detergent and 100 μL of 10% egg yolk phospholipids (w/v) in the form of sonicated liposomes. To this mixture, 50 mM K^+^-gluconate and 20 mM HEPES/Tris (pH 7.0) were added to a final volume of 700 μL. The reconstitution blend was then exposed to 0.5 g of Amberlite XAD-4 resin while continuously stirred at 1200 rev/min at 23 °C for 40 min.

### Transport measurements and transport assay

Following reconstitution, 600 μL of proteoliposomes were loaded onto a Sephadex G-75 column (0.7 cm diameter × 15 cm height) pre-equilibrated with 20 mM HEPES/Tris (pH 7.0) containing 100 mM sucrose to balance the internal osmolarity. Valinomycin (0.75 μg/mg phospholipid) prepared in ethanol was introduced into the eluted proteoliposomes to create a K^+^ diffusion potential. Following a 10 s incubation with valinomycin, transport was initiated by adding 5 μM [^3^H]-Neu5Ac, 0.5 μM *Fn*SiaP, and 10 mM Na^+^-gluconate to 100 μL of liposomes. The initial transport rate was determined by halting the reaction after 15 min, which fell within the initial linear range of [^3^H]-Neu5Ac uptake into the proteoliposomes, as established through the time-course experiments.

The transport assay was concluded by loading each proteoliposome sample (100 μL) onto a Sephadex G-75 column (0.6 cm diameter × 8 cm height) to eliminate external radioactivity. The experimental values were corrected by subtracting the control, i.e. the radioactivity taken up in liposomes reconstituted in the absence of protein. The radioactivity associated with the control samples, i.e. empty liposomes was less than 10% with respect to proteoliposomes. Proteoliposomes were eluted using 1 mL of 50 mM NaCl and the collected eluate was mixed with 4 mL of scintillation mixture, followed by vortexing and counting. Data analysis was conducted using Grafit software (version 5.0.13) using the first-order equation for time-course analysis. As specified in the figure legend, all measurements are presented as mean ± s.d. from at least three independent experiments.

## Supporting information

Combined-supplementary-data-file

## Data availability

The cryo-EM maps and the coordinates of the Neu5Ac bound and the unliganded transporter have been deposited in the EMDB (ID 38926 & 38925), and in the Protein Data Bank (8Y4X & 8Y4W) respectively.

## Acknowledgments

We would like to acknowledge the University of Kentucky’s Nanobody Production Facility for its assistance. We also thank Prof. Jeff Abramson (UCLA) for providing us with the pWARF vector.

## Author contributions

PG expressed and purified all the protein samples for this study. KD was involved in the production of nanobodies and protein purification. With the help of KRV, PG optimized grid preparation and data collection in cryo-EM. PG, RS, and KRV performed the computational work on the cryo-EM data. PG, KRV, and RS designed experiments. MS and CI performed the proteoliposome assays and interpreted the data. RF, RCJD, and RS conceived of the project. All the authors contributed to the writing and revision of the manuscript.

## Funding and additional information

PG, KRV, and SR acknowledge the Department of Biotechnology for the cryo-EM and computing facility, funded under the B-Life grant DBT/PR12422/MED/31/287/2014. SR and RF, acknowledge funding from the Indo-Swedish collaborative grant from the DBT and the Swedish Research Council (BT/IN/Sweden/41/2013). This work was also supported by the DBT/Wellcome Trust India Alliance Fellowship (grant number IA/E/16/1/502999) awarded to PG. R.C.J.D. acknowledges the following for funding support: 1) the Marsden Fund Council from Government funding, managed by Royal Society Te Apārangi (contracts UOC1506 and UOC2211); 2) a Ministry of Business, Innovation, and Employment Smart Ideas grant (contract UOCX1706); and 3) the Biomolecular Interactions Center (UC).

## Ethics Statement

No human subjects were involved in the study.

## Abbreviations

Neu5Ac: *N*-acetylneuraminic acid
TRAP: Tripartite ATP-independent periplasmic
Cryo-EM: Cryo-Electron Microscopy
DDM: n-dodecyl-β-D-maltopyranoside
SiaP: Neu5Ac TRAP substrate-binding protein
SiaQM: Neu5Ac TRAP membrane transporter.

## Supporting information

**Supplementary Figure 1:** Superposition of SiaQM structure from *F. nucleatum*, *H. influenzae* and *P. profundum* with respective bound nanobody or megabody. All the three structures are in inward-open conformation.

**Supplementary Figure 2:** Size exclusion chromatography of TRAP transporter from *Fusobacterium nucleatum* (FnSiaQM). The protein purified in the detergent micelles produced homogenous single peak in size exclusion chromatography eluting as monomer. The purified protein migrated around 50kDa (inset, FnTRAP purified) on a denaturing PAGE, differing from its calculated molecular weight of ∼73.8kDa, which is common for membrane proteins.

**Supplementary Figure 3:** Cryo-EM workflow and analysis of the *Fn*SiaQM protein. **A)** and **B)** a detailed workflow outlining the steps involved in cryo-EM image acquisition and processing, leading to the generation of both the bound and unliganded structures of the *Fn*SiaQM protein, respectively. The selected 2D class average utilized for *ab initio* reconstructions is depicted, and the optimal 3D reconstructions serve as reference models for subsequent non-uniform refinement. Masks generated using RELION 3.0 were applied, and the resulting maps underwent iterative rounds of Non-Uniform refinement, local refinement, 2D classification, selection of 2D classes, and particles representing the selected 2D classes were used for refinement. Final particles were corrected for global and local CTF and reference-based motion correction before final maps were obtained. Fourier shell correlation (FSC) curves for the final 3D reconstructions of the bound and unliganded structures of the *Fn*SiaQM protein are presented. **(C) and (D)** represent the local resolution maps of the unliganded and the liganded *Fn*SiaQM respectively. Much of the interior of the protein including the ligand binding site are well ordered and are the best resolved regions. (E) and (F) Density maps of the unliganded (E) and liganded (F) FnTRAP helices. (G) and (H) The grey surface represents the areas of the density that have been modelled in the unliganded (G) and the liganded (H) forms. In red are the unmodeled densities. The nano-disc surrounding the membrane region is visible as well as number of densities that possibly are lipids bound to the protein. The three figures for each structure represent the different orientations, with the orientation on the right is looking down from the cytosolic side of the protein. The blue density in panel H is the bound sialic acid in the liganded structure.

**Supplementary Figure 4:** SiaQM protein sequence alignment. SiaQM protein sequences from *Photobacterium profundum*, *Haemophilus influenzae,* and *Fusobacterium nucleatum*. Protein sequences were aligned using ESPript 3.

**Supplementary Figure 5:** Ribbon diagram showing the superposition of the *Hi*SiaQM, *Pp*SiaQM and *Fn*SiaQM (liganded and unliganded) structures. The positions of the known Na^+^ ions and metal ion (M) are in spheres, and the position of Neu5Ac is shown in ball and stick.

**Supplementary Figure 6. A)** Na1 site coordination and its interactions with neighboring amino acids; **B)** Na2 site coordination and its interactions with neighboring amino acids; **C)** Metal binding site coordination and its interactions with neighboring amino acids.

**Supplementary Figure 7.** An overlay of sodium ions binding helix-loop-helix regions. In red is SiaQM, and in purple is the structure of VcIndy (PDB ID: 7T9G). The insets display the interactions at the two sodium sites. The red insets show the interactions of SiaQM, while the purple insets represent those of VcIndy.

**Supplementary Figure 8. A and B**. Density of two lipids (Lipid1 and Lipid2) was observed in the scaffold side of *Fn*SiaQM (contour of 1.0 r.m.s. in PyMol). **C.** Electrostatic potential visualization of *Fn*SiaQM. Neu5Ac density in the binding pocket of *Fn*SiaQM. ABPS colored in the range from −5 to +5.

## References

Adams PD, Afonine PV, Bunkoczi G, Chen VB, Davis IW, Echols N, Headd JJ, Hung L-W, Kapral GJ, Grosse-Kunstleve RW, McCoy AJ, Moriarty NW, Oeffner R, Read RJ, Richardson DC, Richardson JS, Terwilliger TC, Zwart PH. 2010. PHENIX: a comprehensive Python-based system for macromolecular structure solution. Acta Crystallographica Section D.

Allen S, Zaleski A, Johnston JW, Gibson BW, Apicella MA. 2005. Novel sialic acid transporter of Haemophilus influenzae. Infection and immunity 73:5291–300. doi:10.1128/IAI.73.9.5291-5300.2005

Almagro-Moreno S, Boyd EF. 2010. Bacterial catabolism of nonulosonic (sialic) acid and fitness in the gut. Gut Microbes 1:45–50. doi:10.4161/gmic.1.1.10386

Bell A, Severi E, Owen CD, Latousakis D, Juge N. 2023. Biochemical and structural basis of sialic acid utilization by gut microbes. Journal of Biological Chemistry 299:102989. doi:10.1016/j.jbc.2023.102989

Bose S, Purkait D, Joseph D, Nayak V, Subramanian R. 2019. Structural and functional characterization of CMP-N-acetylneuraminate synthetase from Vibrio cholerae. Acta Crystallographica Section D: Structural Biology 75:564–577. doi:10.1107/S2059798319006831

Chen X, Varki A. 2010. Advances in the biology and chemistry of sialic acids. ACS chemical biology 5:163–176.

Chou HH, Takematsu H, Diaz S, Iber J, Nickerson E, Wright KL, Muchmore EA, Nelson DL, Warren ST, Varki A. 1998. A mutation in human CMP-sialic acid hydroxylase occurred after the Homo-Pan divergence. Proceedings of the National Academy of Sciences of the United States of America 95:11751–11756. doi:10.1073/pnas.95.20.11751

Chow KM, Whiteheart SW, Smiley JR, Sharma S, Boaz K, Coleman MJ, Maynard A, Hersh LB, Vander Kooi CW. 2019. Immunization of alpacas (Lama pacos) with protein antigens and production of antigen-specific single domain antibodies. JoVE (Journal of Visualized Experiments) e58471.

Coombes D, Davies JS, Newton-Vesty MC, Horne CR, Setty TG, Subramanian R, Moir JWB, Rosmarie Friemann X, Panjikar S, Griffin MDW, North RA, Dobson RCJ. 2020. The basis for non-canonical ROK family function in the N-acetylmannosamine kinase from the pathogen Staphylococcus aureus. Journal of Biological Chemistry 295:3301–3315. doi:10.1074/jbc.RA119.010526

Corfield AP. 2015. Mucins: A biologically relevant glycan barrier in mucosal protection. Biochimica et Biophysica Acta (BBA) - General Subjects 1850:236–252. doi:10.1016/j.bbagen.2014.05.003

Currie MJ, Davies JS, Scalise M, Gulati A, Wright JD, Newton-Vesty MC, Abeysekera GS, Subramanian R, Wahlgren WY, Friemann R, Allison JR, Mace PD, Griffin MD, Demeler B, Wakatsuki S, Drew D, Indiveri C, Dobson RC, North RA. 2024. Structural and biophysical analysis of a Haemophilus influenzae tripartite ATP-independent periplasmic (TRAP) transporter. eLife 12:RP92307. doi:10.7554/eLife.92307

Currie MJ, Manjunath L, Horne CR, Rendle PM, Subramanian R, Friemann R, Fairbanks AJ, Muscroft-Taylor AC, North RA, Dobson RCJ. 2021. N-acetylmannosamine-6-phosphate 2-epimerase uses a novel substrate-assisted mechanism to catalyze amino sugar epimerization. Journal of Biological Chemistry 297. doi:10.1016/j.jbc.2021.101113

Davies JS, Coombes D, Horne CR, Pearce FG, Friemann R, North RA, Dobson RC. 2019. Functional and solution structure studies of amino sugar deacetylase and deaminase enzymes from Staphylococcus aureus. FEBS letters 593:52–66.

Davies JS, Currie MJ, North RA, Scalise M, Wright JD, Copping JM, Remus DM, Gulati A, Morado DR, Jamieson SA, Newton-Vesty MC, Abeysekera GS, Ramaswamy S, Friemann R, Wakatsuki S, Allison JR, Indiveri C, Drew D, Mace PD, Dobson RCJ. 2023. Structure and mechanism of a tripartite ATP-independent periplasmic TRAP transporter. Nat Commun 14:1120. doi:10.1038/s41467-023-36590-1

DeLano WL. 2002. The PyMOL molecular graphics system. http://www.pymol.org/.

Denisov IG, Sligar SG. 2016. Nanodiscs for structural and functional studies of membrane proteins. Nat Struct Mol Biol 23:481–486. doi:10.1038/nsmb.3195

Emsley P, Cowtan K. 2004. Coot: model-building tools for molecular graphics. Acta crystallographica section D: biological crystallography 60:2126–2132.

Evans R, O’Neill M, Pritzel A, Antropova N, Senior A, Green T, Žídek A, Bates R, Blackwell S, Yim J, Ronneberger O, Bodenstein S, Zielinski M, Bridgland A, Potapenko A, Cowie A, Tunyasuvunakool K, Jain R, Clancy E, Kohli P, Jumper J, Hassabis D. 2022. Protein complex prediction with AlphaFold-Multimer. doi:10.1101/2021.10.04.463034

Felder CB, Graul RC, Lee AY, Merkle H-P, Sadee W. 1999. The venus flytrap of periplasmic binding proteins: An ancient protein module present in multiple drug receptors. AAPS PharmSci 1:7–26. doi:10.1208/ps010202

Gangi Setty T, Cho C, Govindappa S, Apicella MA, Ramaswamy S. 2014. Bacterial periplasmic sialic acid-binding proteins exhibit a conserved binding site. Acta Cryst D 70:1801–1811. doi:10.1107/S139900471400830X

Garaeva AA, Guskov A, Slotboom DJ, Paulino C. 2019. A one-gate elevator mechanism for the human neutral amino acid transporter ASCT2. Nat Commun 10:3427. doi:10.1038/s41467-019-11363-x

Glaenzer J, Peter MF, Thomas GH, Hagelueken G. 2017. PELDOR Spectroscopy Reveals Two Defined States of a Sialic Acid TRAP Transporter SBP in Solution. Biophys J 112:109–120. doi:10.1016/j.bpj.2016.12.010

Glanz VY, Myasoedova VA, Grechko AV, Orekhov AN. 2018. Inhibition of sialidase activity as a therapeutic approach. Drug Des Devel Ther 12:3431–3437. doi:10.2147/DDDT.S176220

Haines-Menges BL, Whitaker WB, Lubin JB, Boyd EF. 2015. Host Sialic Acids: A Delicacy for the Pathogen with Discerning Taste. Metabolism and Bacterial Pathogenesis 3:321–342. doi:10.1128/microbiolspec.MBP-0005-2014

Horne CR, Kind L, Davies JS, Dobson RC. 2020. On the structure and function of Escherichia coli YjhC: an oxidoreductase involved in bacterial sialic acid metabolism. Proteins: Structure, Function, and Bioinformatics 88:654–668.

Hsieh JM, Besserer GM, Madej MG, Bui H-Q, Kwon S, Abramson J. 2010. Bridging the gap: a GFP-based strategy for overexpression and purification of membrane proteins with intra and extracellular C-termini. Protein Science 19:868–880.

Johnston JW, Coussens NP, Allen S, Houtman JCD, Turner KH, Zaleski A, Ramaswamy S, Gibson BW, Apicella MA. 2008. Characterization of the N-Acetyl-5-neuraminic Acid-binding Site of the Extracytoplasmic Solute Receptor (SiaP) of Nontypeable Haemophilus influenzae Strain 2019 *. Journal of Biological Chemistry 283:855–865. doi:10.1074/jbc.M706603200

Jurrus E, Engel D, Star K, Monson K, Brandi J, Felberg LE, Brookes DH, Wilson L, Chen J, Liles K, Chun M, Li P, Gohara DW, Dolinsky T, Konecny R, Koes DR, Nielsen JE, Head-Gordon T, Geng W, Krasny R, Wei G-W, Holst MJ, McCammon JA, Baker NA. 2018. Improvements to the APBS biomolecular solvation software suite. Protein Science 27:112–128. doi:10.1002/pro.3280

Kawate T, Gouaux E. 2006. Fluorescence-detection size-exclusion chromatography for precrystallization screening of integral membrane proteins. Structure (London, England : 1993) 14:673–81. doi:10.1016/j.str.2006.01.013

Kinz-Thompson CD, Lopez-Redondo ML, Mulligan C, Sauer DB, Marden JJ, Song J, Tajkhorshid E, Hunt JF, Stokes DL, Mindell JA, Wang DN, Gonzalez RL. 2022. Elevator mechanism dynamics in a sodium-coupled dicarboxylate transporter. doi:10.1101/2022.05.01.490196

Kumar JP, Rao H, Nayak V, Ramaswamy S. 2018. Crystal structures and kinetics of N-acetylneuraminate lyase from Fusobacterium nucleatum. Acta Crystallographica Section F: Structural Biology Communications 74:725–732. doi:10.1107/S2053230X18012992

Lewis AL, Lewis WG. 2012. Host sialoglycans and bacterial sialidases: a mucosal perspective. Cellular Microbiology 14:1174–1182. doi:10.1111/j.1462-5822.2012.01807.x

Mancusso R, Gregorio GG, Liu Q, Wang D-N. 2012. Structure and mechanism of a bacterial sodium-dependent dicarboxylate transporter. Nature 491:622–626. doi:10.1038/nature11542

Manjunath L, Guntupalli SR, Currie MJ, North RA, Dobson RCJ, Nayak V, Subramanian R. 2018. Crystal structures and kinetic analyses of *N*-acetylmannosamine-6-phosphate 2-epimerases from *Fusobacterium nucleatum* and *Vibrio cholerae*. Acta Crystallographica Section F Structural Biology Communications 74:431–440. doi:10.1107/S2053230X18008543

Müller A, Severi E, Mulligan C, Watts AG, Kelly DJ, Wilson KS, Wilkinson AJ, Thomas GH. 2006. Conservation of Structure and Mechanism in Primary and Secondary Transporters Exemplified by SiaP, a Sialic Acid Binding Virulence Factor from Haemophilus influenzae*. Journal of Biological Chemistry 281:22212–22222. doi:10.1074/jbc.M603463200

Mulligan C, Geertsma ER, Severi E, Kelly DJ, Poolman B, Thomas GH. 2009. The substrate-binding protein imposes directionality on an electrochemical sodium gradient-driven TRAP transporter. Proceedings of the National Academy of Sciences of the United States of America 106:1778–83. doi:10.1073/pnas.0809979106

Mulligan C, Leech AP, Kelly DJ, Thomas GH. 2012. The membrane proteins SiaQ and SiaM form an essential stoichiometric complex in the sialic acid tripartite ATP-independent periplasmic (TRAP) transporter SiaPQM (VC1777-1779) from Vibrio cholerae. The Journal of biological chemistry 287:3598–3608. doi:10.1074/jbc.M111.281030

Nie R, Stark S, Symersky J, Kaplan RS, Lu M. 2017. Structure and function of the divalent anion/Na+ symporter from Vibrio cholerae and a humanized variant. Nat Commun 8:15009. doi:10.1038/ncomms15009

North RA, Wahlgren WY, Remus DM, Scalise M, Kessans SA, Dunevall E, Claesson E, Soares da Costa TP, Perugini MA, Ramaswamy S, Allison JR, Indiveri C, Friemann R, Dobson RCJ. 2018. The Sodium Sialic Acid Symporter From Staphylococcus aureus Has Altered Substrate Specificity. Frontiers in Chemistry 6:233. doi:10.3389/fchem.2018.00233

North RA, Watson AJ, Pearce FG, Muscroft-Taylor AC, Friemann R, Fairbanks AJ, Dobson RC. 2016. Structure and inhibition of N-acetylneuraminate lyase from methicillin-resistant Staphylococcus aureus. FEBS letters 590:4414–4428.

Peter MF, Gebhardt C, Glaenzer J, Schneberger N, de Boer M, Thomas GH, Cordes T, Hagelueken G. 2021. Triggering Closure of a Sialic Acid TRAP Transporter Substrate Binding Protein through Binding of Natural or Artificial Substrates. Journal of Molecular Biology 433:166756. doi:10.1016/j.jmb.2020.166756

Peter MF, Ruland JA, Depping P, Schneberger N, Severi E, Moecking J, Gatterdam K, Tindall S, Durand A, Heinz V, Siebrasse JP, Koenig P-A, Geyer M, Ziegler C, Kubitscheck U, Thomas GH, Hagelueken G. 2022. Structural and mechanistic analysis of a tripartite ATP-independent periplasmic TRAP transporter. Nat Commun 13:4471. doi:10.1038/s41467-022-31907-y

Peter MF, Ruland JA, Kim Y, Hendricks P, Schneberger N, Siebrasse JP, Thomas GH, Kubitscheck U, Hagelueken G. 2024. Conformational coupling of the sialic acid TRAP transporter HiSiaQM with its substrate binding protein HiSiaP. Nat Commun 15:217. doi:10.1038/s41467-023-44327-3

Pettersen EF, Goddard TD, Huang CC, Couch GS, Greenblatt DM, Meng EC, Ferrin TE. 2004. UCSF Chimera—a visualization system for exploratory research and analysis. Journal of computational chemistry 25:1605–1612.

Punjani A. 2020. Algorithmic Advances in Single Particle Cryo-EM Data Processing Using CryoSPARC. Microscopy and Microanalysis 26:2322–2323. doi:10.1017/S1431927620021194

Punjani A, Rubinstein JL, Fleet DJ, Brubaker MA. 2017. cryoSPARC: algorithms for rapid unsupervised cryo-EM structure determination. Nature Methods 14:290–296. doi:10.1038/nmeth.4169

Rosa LT, Bianconi ME, Thomas GH, Kelly DJ. 2018. Tripartite ATP-Independent Periplasmic (TRAP) Transporters and Tripartite Tricarboxylate Transporters (TTT): From Uptake to Pathogenicity. Front Cell Infect Microbiol 8:33. doi:10.3389/fcimb.2018.00033

Sauer DB, Marden JJ, Sudar JC, Song J, Mulligan C, Wang D-N. 2022. Structural basis of ion – substrate coupling in the Na+-dependent dicarboxylate transporter VcINDY. Nat Commun 13:2644. doi:10.1038/s41467-022-30406-4

Setty TG, Mowers JC, Hobbs AG, Maiya SP, Syed S, Munson RS, Apicella MA, Subramanian R. 2018. Molecular characterization of the interaction of sialic acid with the periplasmic binding protein from Haemophilus ducreyi. Journal of Biological Chemistry 293:20073–20084. doi:10.1074/jbc.RA118.005151

Severi E, Hood DW, Thomas GH. 2007. Sialic acid utilization by bacterial pathogens. Microbiology 153:2817–2822. doi:10.1099/mic.0.2007/009480-0

Sillanaukee, Pönniö, Jääskeläinen. 1999. Occurrence of sialic acids in healthy humans and different disorders. European Journal of Clinical Investigation 29:413–425. doi:10.1046/j.1365-2362.1999.00485.x

Varki A. 2009. Multiple changes in sialic acid biology during human evolution. Glycoconj J 26:231–245. doi:10.1007/s10719-008-9183-z

Vetting MW, Al-Obaidi N, Zhao S, San Francisco B, Kim J, Wichelecki DJ, Bouvier JT, Solbiati JO, Vu H, Zhang X, Rodionov DA, Love JD, Hillerich BS, Seidel RD, Quinn RJ, Osterman AL, Cronan JE, Jacobson MP, Gerlt JA, Almo SC. 2015. Experimental Strategies for Functional Annotation and Metabolism Discovery: Targeted Screening of Solute Binding Proteins and Unbiased Panning of Metabolomes. Biochemistry 54:909–931. doi:10.1021/bi501388y

Wahlgren WY, Dunevall E, North RA, Paz A, Scalise M, Bisignano P, Bengtsson-Palme J, Goyal P, Claesson E, Caing-Carlsson R, Andersson R, Beis K, Nilsson UJ, Farewell A, Pochini L, Indiveri C, Grabe M, Dobson RCJ, Abramson J, Ramaswamy S, Friemann R. 2018. Substrate-bound outward-open structure of a Na+-coupled sialic acid symporter reveals a new Na+site. Nature Communications 9:1753. doi:10.1038/s41467-018-04045-7

Wang S, Xie J, Pei J, Lai L. 2023. CavityPlus 2022 Update: An Integrated Platform for Comprehensive Protein Cavity Detection and Property Analyses with User-friendly Tools and Cavity Databases. Journal of Molecular Biology, Computation Resources for Molecular Biology 435:168141. doi:10.1016/j.jmb.2023.168141

Wang X, Boudker O. 2020. Large domain movements through the lipid bilayer mediate substrate release and inhibition of glutamate transporters. eLife 9:e58417. doi:10.7554/eLife.58417

Wirth C, Condemine G, Boiteux C, Bernèche S, Schirmer T, Peneff CM. 2009. NanC Crystal Structure, a Model for Outer-Membrane Channels of the Acidic Sugar-Specific KdgM Porin Family. Journal of Molecular Biology 394:718–731. doi:10.1016/j.jmb.2009.09.054

