## Supplementary material for "Molecular determinants of Neu5Ac binding to a tripartite ATP independent periplasmic (TRAP) transporter": Combined-supplementary-data-file

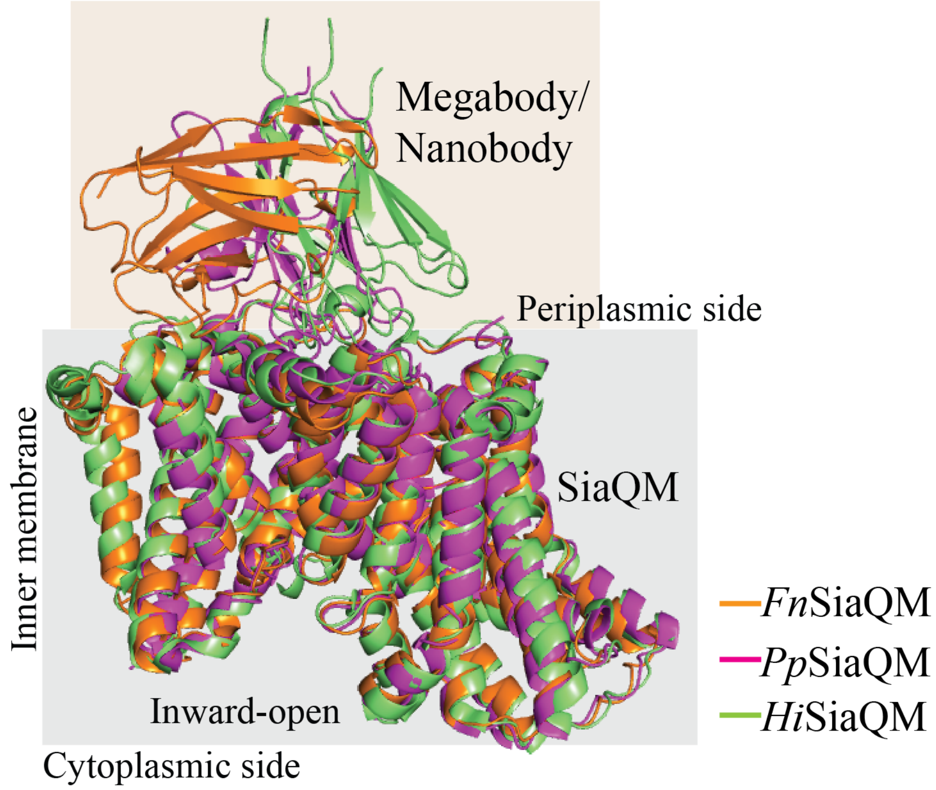


**Supplementary Figure 1.** Superposition of SiaQM structure from *F. nucleatum*, *H. influenzae* and *P. profundum* with respective bound nanobody or megabody. All three structures structures are in inward-open conformation.

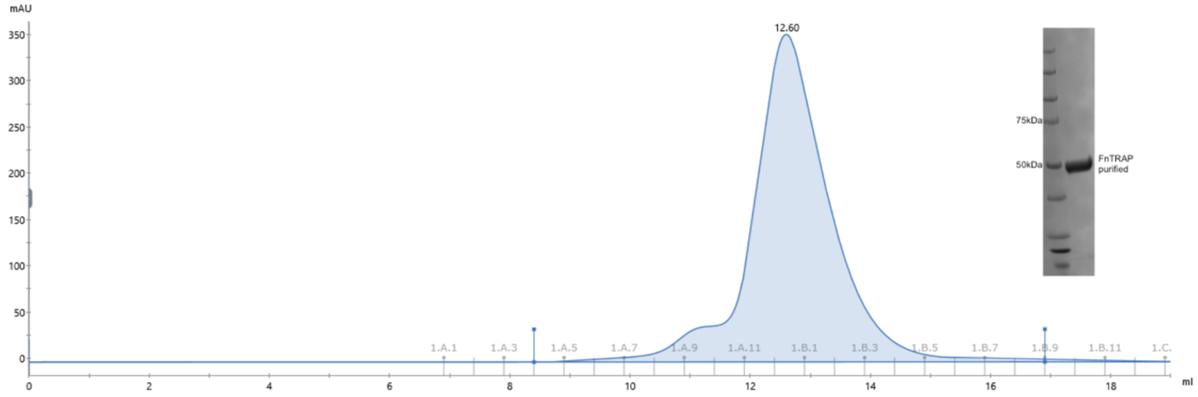


**Supplementary Figure 2.** Size exclusion chromatography of TRAP transporter from *Fusobacterium nucleatum* (FnSiaQM). The protein purified in the detergent micelles produced a homogenous single peak in size exclusion chromatography, eluting as a monomer. The purified protein migrated around 50kDa (inset, FnTRAP purified) on a denaturing PAGE, differing from its calculated molecular weight of ~73.8kDa, which is common for membrane proteins.

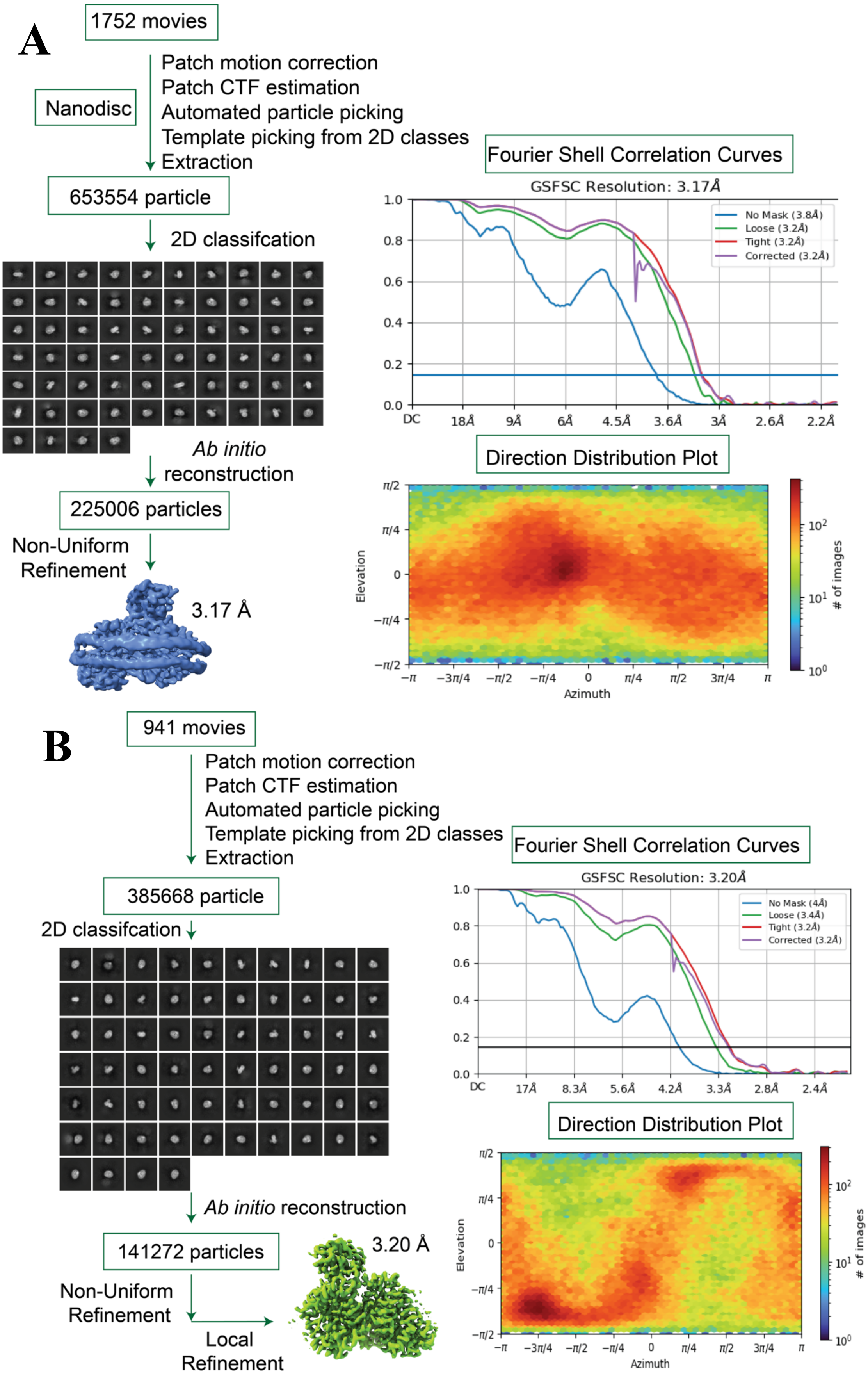
A
1752 movies
Nanodisc
Patch motion correction
Patch CTF estimation
Automated particle picking
Template picking from 2D classes
Extraction
653554 particle
2D classifcation
Ab initio reconstruction
225006 particles
Non-Uniform Refinement
3.17 Å
Fourier Shell Correlation Curves
GSFSC Resolution: 3.17Å
No Mask (3.8Å)
Loose (3.2Å)
Tight (3.2Å)
Corrected (3.2Å)
DC
18Å
9Å
6Å
4.5Å
3.6Å
3Å
2.6Å
2.2Å
Direction Distribution Plot
Elevation
Azimuth
### of images
B
941 movies
Patch motion correction
Patch CTF estimation
Automated particle picking
Template picking from 2D classes
Extraction
385668 particle
2D classifcation
Ab initio reconstruction
141272 particles
Non-Uniform Refinement
Local Refinement
3.20 Å
Fourier Shell Correlation Curves
GSFSC Resolution: 3.20Å
No Mask (4Å)
Loose (3.4Å)
Tight (3.2Å)
Corrected (3.2Å)
DC
17Å
8.3Å
5.6Å
4.2Å
3.3Å
2.8Å
2.4Å
Direction Distribution Plot
Elevation
Azimuth
### of images

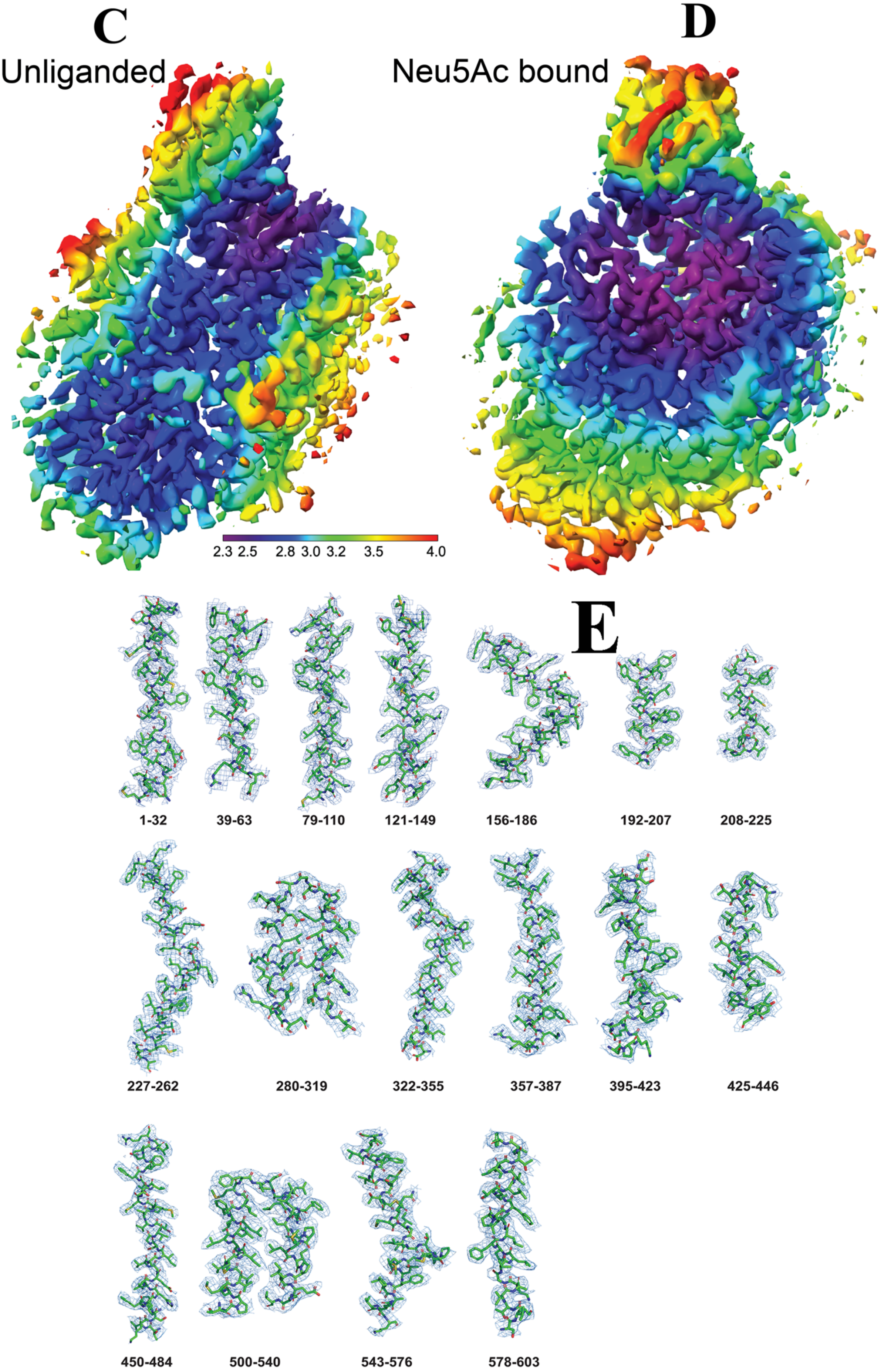
C
Unliganded
D
Neu5Ac bound
2.3 2.5 2.8 3.0 3.2 3.5 4.0
E
1-32
39-63
79-110
121-149
156-186
192-207
208-225
227-262
280-319
322-355
357-387
395-423
425-446
450-484
500-540
543-576
578-603

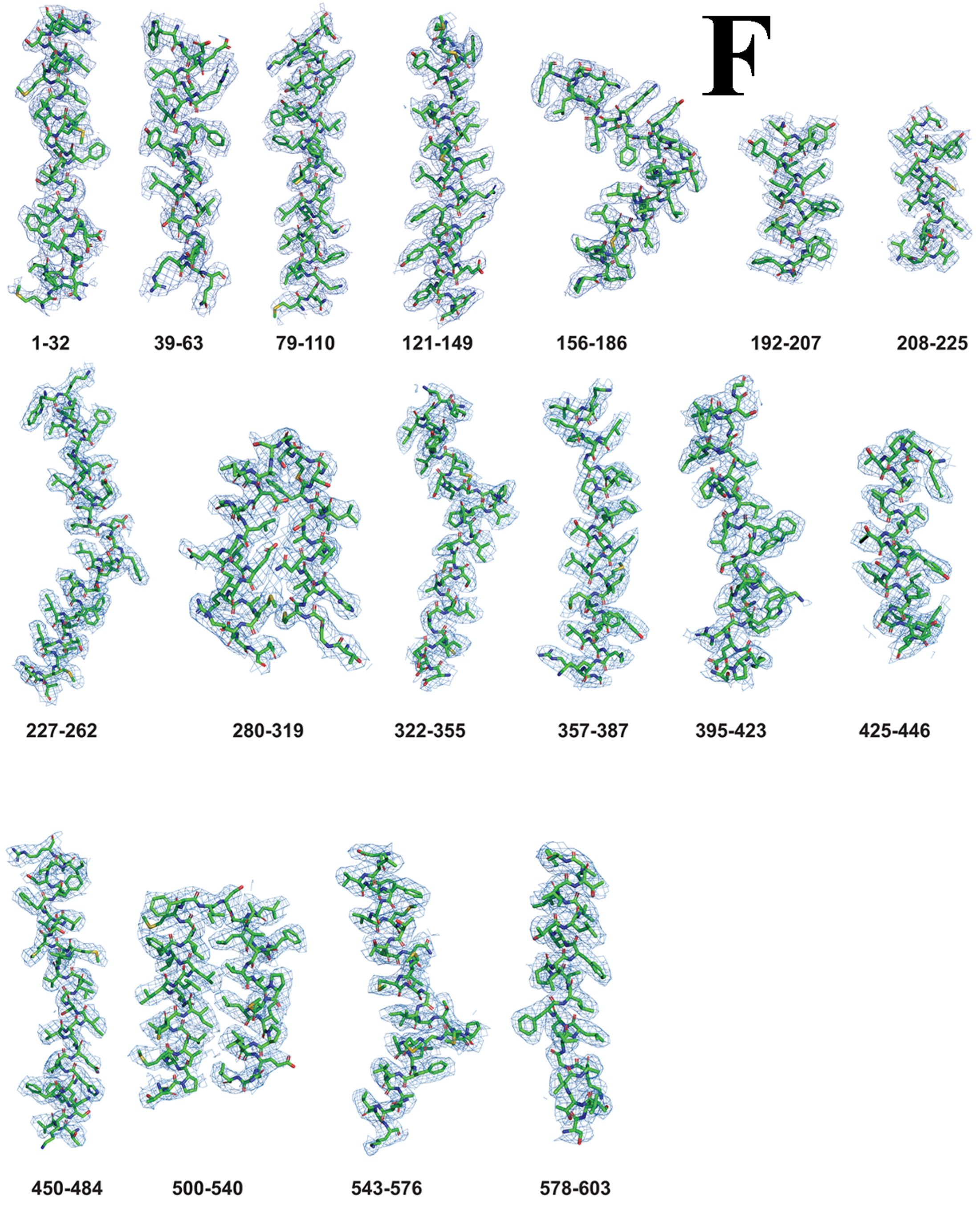
F
1-32
39-63
79-110
121-149
156-186
192-207
208-225
227-262
280-319
322-355
357-387
395-423
425-446
450-484
500-540
543-576
578-603

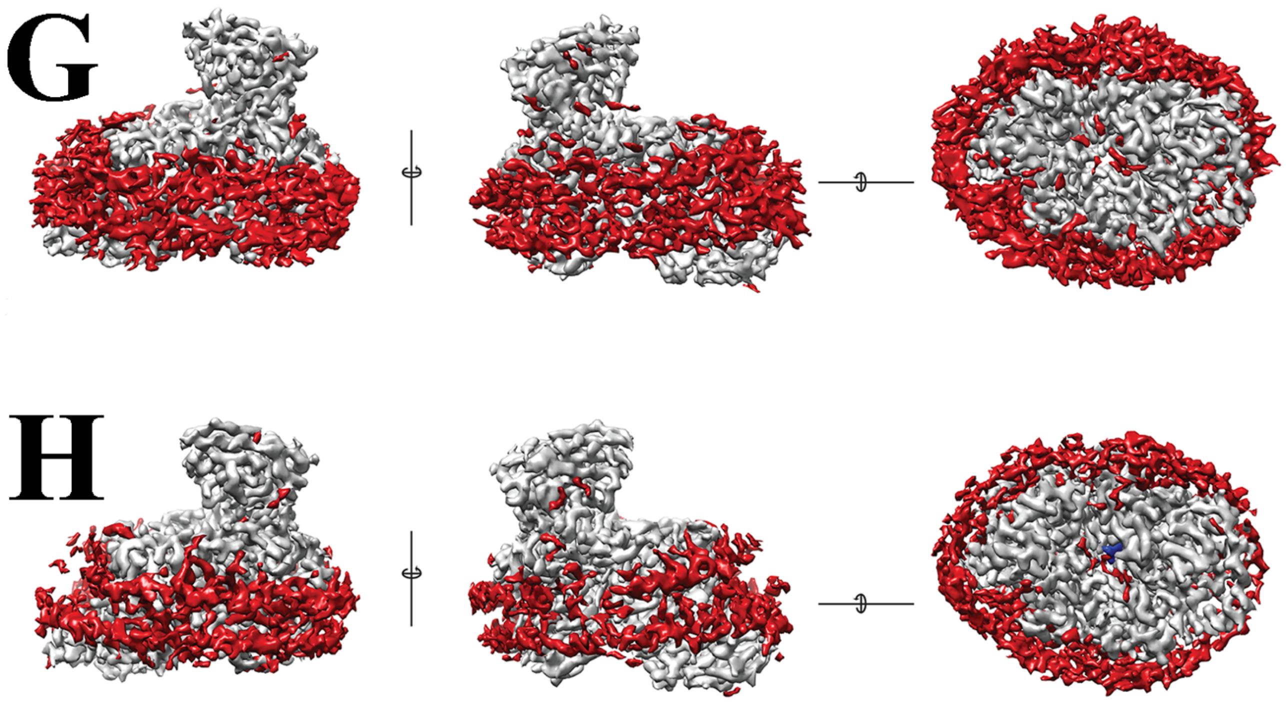


**Supplementary Figure 3.** Cryo-EM workflow and analysis of the *Fn*SiaQM protein. **A) and B)** a detailed workflow outlining the steps involved in cryo-EM image acquisition and processing, leading to the generation of both the bound and unliganded structures of the *Fn*SiaQM protein, respectively. The selected 2D class averages utilized for *ab initio* reconstructions are depicted, and the optimal 3D reconstructions serve as reference models for subsequent non-uniform refinement. Masks generated using RELION 3.0 were applied, and the resulting maps underwent iterative rounds of Non-Uniform refinement, local refinement, 2D classification, selection of 2D classes, and particles representing the selected 2D classes were used for refinement. Final particles were corrected for global and local CTF and reference-based motion correction before final maps were obtained. Fourier shell correlation (FSC) curves for the final 3D reconstructions of the bound and unliganded structures of the *Fn*SiaQM protein are presented. **(C) and (D)** represent the local resolution maps of the unliganded and the liganded *Fn*SiqQM respectively. Most of the interior of the protein including the ligand binding site are well ordered and are the best resolved regions. **(E) and (F)** Density maps of the unliganded (E) and liganded (F) FnTRAP helices. **(G) and (H)** The grey surface represents the areas of the density that have been modelled in the unliganded (G) and the liganded (H) forms. In red are the unmodeled densities. The nano-disc surrounding the membrane region is visible as well as number of densities that possibly are lipids bound to the protein. The three figures for each structure represent the different orientations, with the orientation on the right is looking down from the cytosolic side of the protein. The blue density in panel H is the bound sialic acid in the liganded structure.

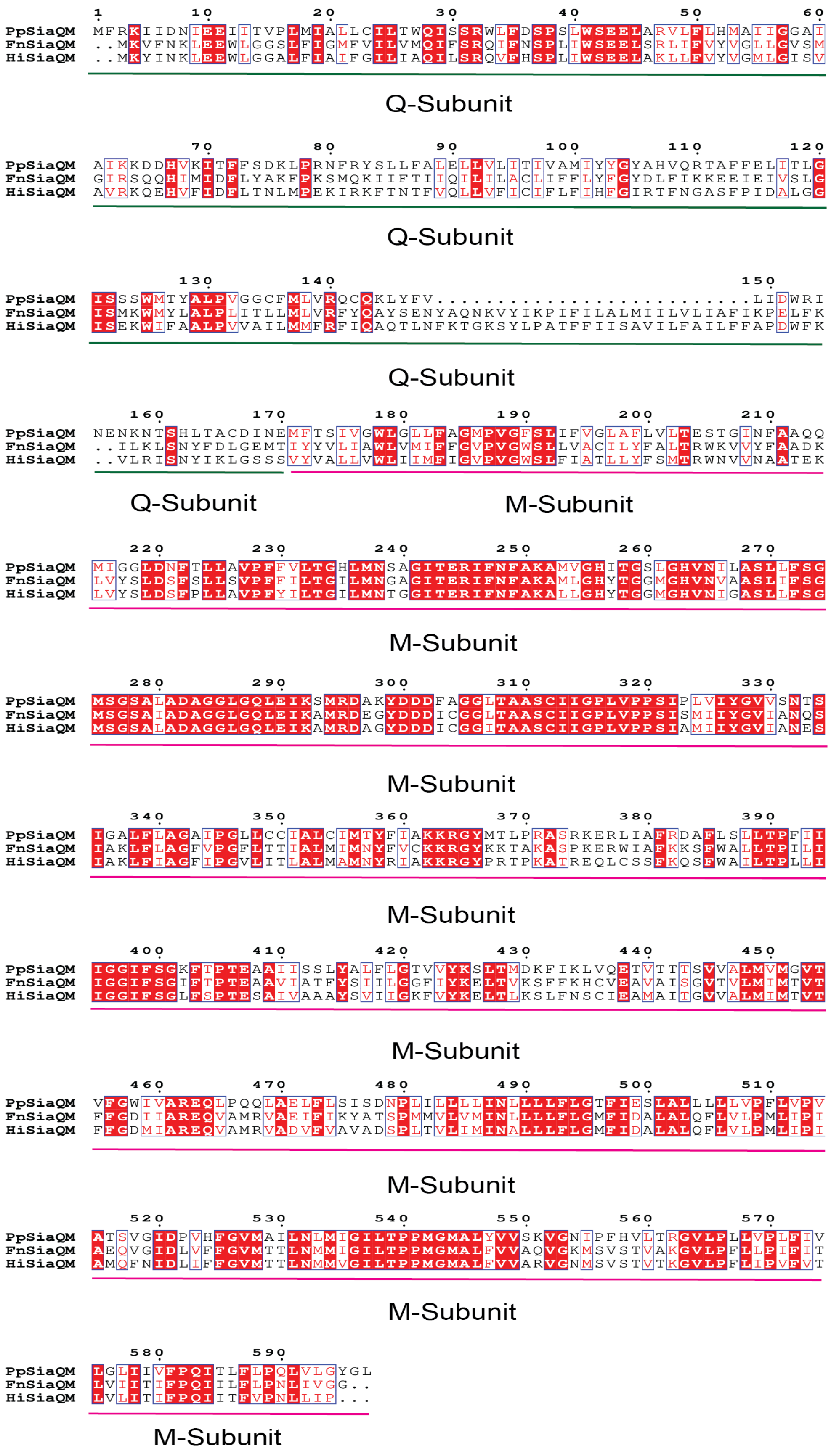


**Supplementary Figure 4.** SiaQM protein sequence alignment. SiaQM protein sequences from *Photobacterium profundum*, *Haemophilus influenzae*, and *Fusobacterium nucleatum*. Protein sequences were aligned using ESPript 3.

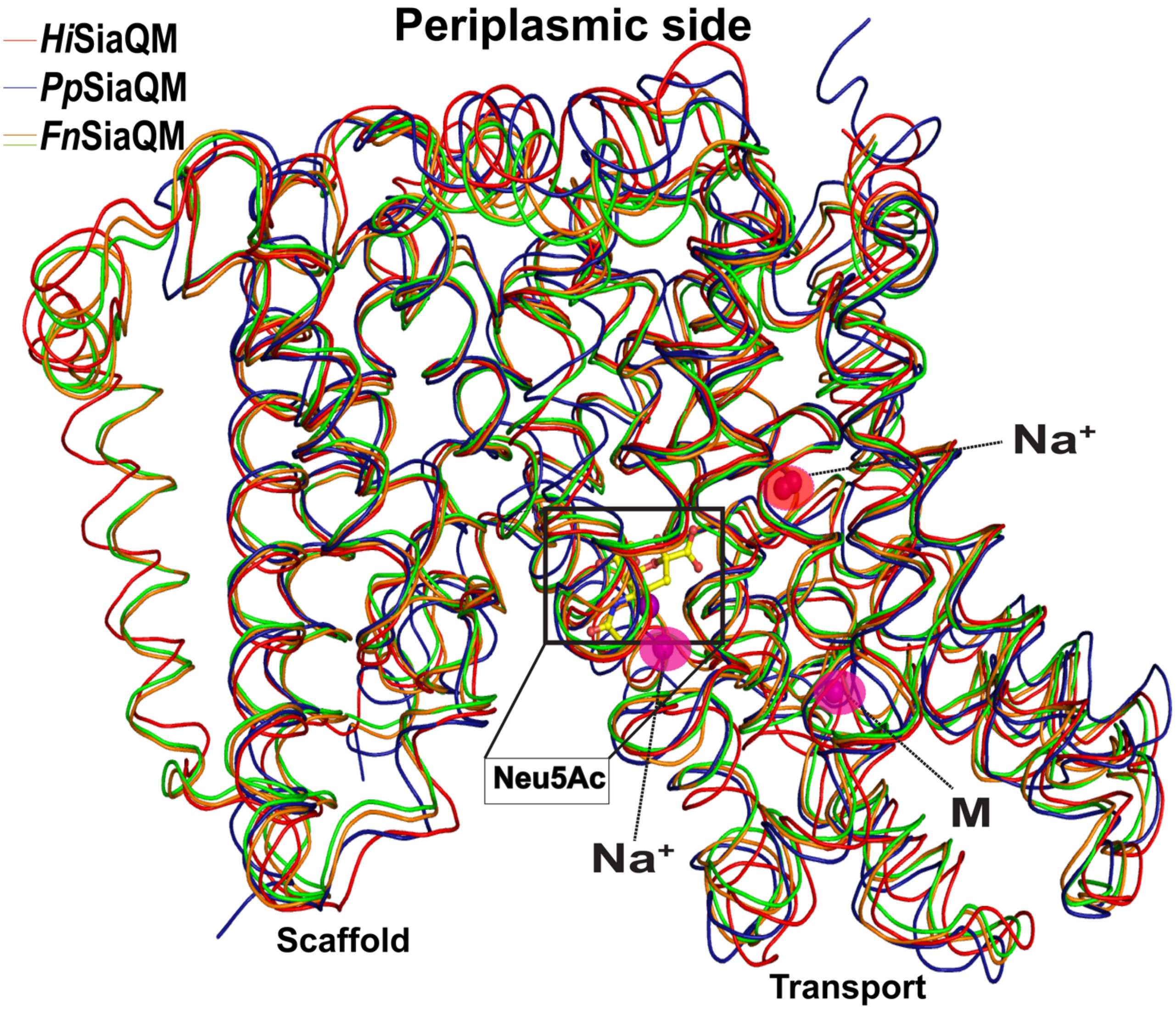


**Supplementary Figure 5.** Ribbon diagram showing the superposition of the *Hi*SiaQM, *Pp*SiaQM and *Fn*SiaQM (liganded and unliganded) structures. The positions of the known $Na^{+}$ ions and metal ion (M) are in spheres, and the position of Neu5Ac is shown in ball and stick.

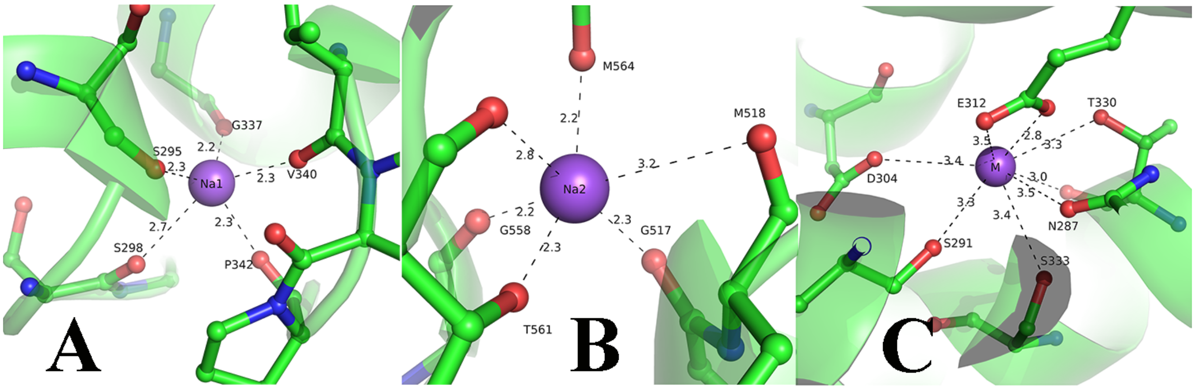


**Supplementary Figure 6. A)** Na1 site coordination and its interactions with neighboring amino acids; **B)** Na2 site coordination and its interactions with neighboring amino acids; **C)** Metal binding site coordination and its interactions with neighboring amino acids.

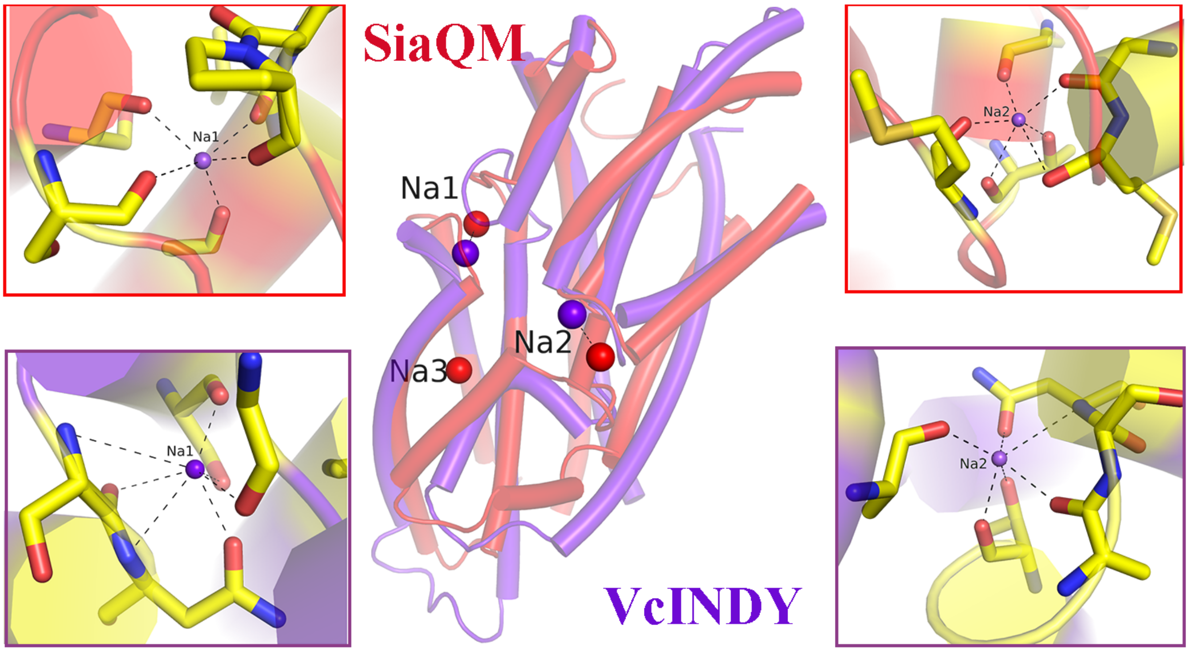


**Supplementary Figure 7.** An overlay of sodium ions binding helix-loop-helix regions. In red is SiaQM, and in purple is the structure of VcIndy (PDB ID: 7T9G). The insets display the interactions at the two sodium sites. The red insets show the interactions of SiaQM, while the purple insets represent those of VcIndy.

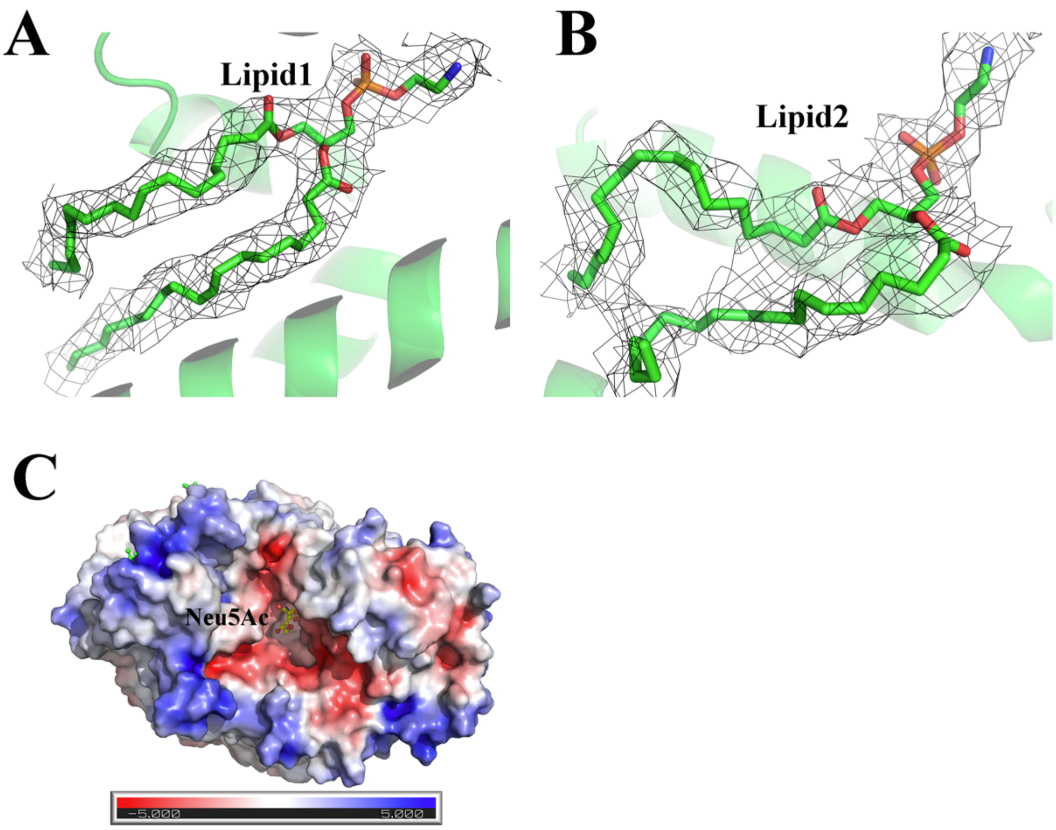


**Supplementary Figure 8. A and B**. Density of two lipids (Lipid1 and Lipid2) was observed in the scaffold side of *Fn*SiaQM (contour of 1.0 r.m.s. in PyMol). **C**. Electrostatic potential visualization of *Fn*SiaQM. Neu5Ac density in the binding pocket of *Fn*SiaQM. ABPS colored in the range from -5 to +5.
